# Transmembrane region dimer structures of Type 1 receptors readily sample alternate configurations: MD simulations using the Martini 3 coarse grained model compared to AlphaFold2 Multimer

**DOI:** 10.1101/2021.09.10.459840

**Authors:** Amita R. Sahoo, Paulo C. T. Souza, Zhiyuan Meng, Matthias Buck

**Author notes:** Biophysics Graduate Program, The Ohio State University, Columbus, Ohio 43210, U. S. A.

## Abstract

Determination of the structure and dynamics of transmembrane (TM) regions of single-transmembrane receptors is key to understanding their mechanism of signal transduction across the plasma membrane. Although many studies have been performed on isolated soluble extra- and intracellular receptor domains in aqueous solutions, limited knowledge exists on the lipid embedded TM domain. In this study, we examine the assembly of configurations of receptor TM region dimers using the Martini 3 force field for coarse-grain (CG) molecular dynamics simulations. This recently published version of Martini has new bead types and sizes, which allows more accurate predictions of molecular interactions compared to the previous versions. At first glance our results with Martini 3 simulations show only a reasonable agreement with *ab initio* predictions using PREDDIMER (for TM domains only), AlphaFold2 Multimer and with available NMR derived structures for TM helix dimers. Specifically, 6 of 11 CG TM structures are similar to the NMR structures (within < 3.5 Å mainchain RMSD) compared to 10 of 11 and 9 of 11 using PREDDIMER and Alphafold, respectively (7 structures of the latter are within 1.5 Å) Surprisingly, AlphaFold2 predictions are more comparable with NMR structures when the database of 2001 (mainly composed of soluble proteins) instead of 2020 PDB structures are used. While there are some differences in the conditions used, the CG simulations primarily reveal that alternate configurations of the TM dimers that are sampled, which readily interconvert with a predominant population. The implications of these findings for our understanding of the signalling mechanism of TM receptors are discussed, including opportunities for the development of new pharmaceuticals, some of which are peptide based.

## Introduction

Membrane proteins account for 20-30 % of all proteins identified in the genomes of prokaryotes and eukaryotes.^1^ Compared with the multi-pass membrane proteins, single-pass transmembrane (type 1) receptors are the most abundant and functionally diverse category of membrane proteins.^2^ These proteins are often highly flexible near the membrane and very difficult to characterize structurally. Signal transduction across the plasma membrane typically involves receptor dimerization, not just of TM regions but also of TM adjacent regions. The TM region, comprising the membrane embedded TM domain and the membrane surrounding region of amino acids, can contribute to the stability of full-length receptor dimers and hence, help maintain configurations which are either competent for signalling or inactive. Specific sets of interhelix contacts between the TM region, set up distinct “on”/ “off” states through change in the orientation, if not oligomerization of the TM regions. It is thought that within a biological membrane, individual TM helices usually interact to form one or only a few thermodynamically stable structures. Properties of the lipid bilayer, such as the thickness of the membrane, the nature of lipid tails and headgroups are also key contributing factors to the stability of TM domain configurations.^3–5^ Solution NMR has been the main tool for the determination of TM helical dimer structures using detergent micelles (typically DPC), or more realistic membrane mimics, such as bicelles, a mixture of DMPC lipid and DHPC detergent. However, the NMR data are typically collected and analysed in a way to determine the structure for one particular configuration of a dimer promoted by such environment^6^ and it is noticeable that most structures were determined at an unphysiologically low pH (Table S1) which tends to stabilize a particular configuration of the TM dimer. In fact, there is a His and/or Glu residue within 6 residues (potentially <2 helical turns, but often closer) to the N- and/or C-terminus of the non-polar TM segment in many (type 1) receptors.

To date the structures of several TM helix homo- or heterodimers have been solved. Analyzing these structures and how they assemble by studying the dynamics of the simple TM helix dimers helps us to understand the type and mode of interactions between individual TM helices, at the level of amino acids and/or amino acid motifs^7^ In this study, we focused on a computational characterization of the structural dynamics of the TM domains of a total of 11 receptors: Bnip3, EphA1, EphA2, ErbB1, ErbB1/2, ErbB2, ErbB3, ErbB4, FEFR3, GpA and PDGFRb.^8–18^ Of the 11 TM regions studied, all are from receptor tyrosine kinases (RTK) with an exception of Bnip3, which is a bcl2-family (non-kinase receptor) and GpA, Glycophorin A, a previously established model system for TM dimers. *Ab initio* predictions have been made for these receptors^19,20^ and further structural refinements using µs level all-atom molecular dynamics (MD) simulations^20,21^ and also CG simulations using the Martini 2 force field have been carried out for some of the 11 systems.^22–25^ Much of the attention has been focused on reproducing the experimental NMR structures. However, the configurational dynamics of the slightly wider TM region at neutral pH remain far less explored and this is our focus here, given that Martini 3^26^ has been reasonably validated with several examples of TM region helix dimers.

All-atom simulations have been employed to study the binding mode of TM helices previously, but since the movement of lipids and TM helices in lipids is relatively slow, these methods demand extensive computational resources that are difficult to access.^27,28^ For example, in the landmark 2013 study of EGFR TM by Kuriyan and Shaw,^29^ it took > 100 μs for the TM regions to dimerize correctly. Significant computational speed-ups (typically 100 to 1000-fold over all-atom MD) can be achieved by carrying out coarse grained (CG) simulations using a recently developed Martini 3 force field.^26^ This version has new particles with improved packing and interaction balance of all the non-bonded interaction terms, allowing more realistic modelling of biomolecular systems, including protein-protein and protein-membrane interactions examined here. Using Martini 3, we gained insights into how these 11 TM peptides associate/dimerize in the CG simulations in a DMPC lipid bilayer. Surprisingly we obtained a predominant structural ensemble, but also several side, alternate configurational states in most cases. Significantly, these structures are seen to readily interconvert in interhelix crossing angle, even when the helices only modestly separated. Comparison with *ab initio* PREDDIMER,^19^ Alphafold2 Multimer^30^ predictions and the solution NMR structures^8–18^ suggest that the presence of alternate structures and their interconversion in the CG simulations likely arises, at least in part, from regions just outside the TM hydrophobic domain. This view is consistent with experimental findings and emphasizes the multistate nature of transmembrane proteins, where further extra- and intracellular domain interactions likely synergize to define a few TM crossing configurational states.

## Results and Discussion

The non-polar segment, that is membrane-crossing part of the transmembrane (TM) region for the 11 receptors were modeled as ideal α-helices. Because the regions immediately outside this TM segment can affect the structure of the TM region and stability of the dimeric configuration states, we extended the N-terminus and C-terminus of the TM region by 8-9 residues and 10 residues, respectively (Table S2), modelling them initially as extended conformation. For each protein, we modeled two identical TM peptides and placed them in parallel to the membrane normal, in the center of the DMPC bilayer with an interhelical separation of 50 Å from each other (Fig. 1). It is known that the configuration of the TM helices can be affected by membrane thickness^31^ but also by lipid composition, e.g. presence of cholesterol.^32,33^ However, in order to best compare with the experimental data on the NMR structures in DMPC/DHPC bicelles and some in DPC, we chose the same DMPC bilayer throughout. Each system was then run in quadruplicate for 4 µs each. The TM peptides came closer via diffusion in the membrane, and typically within 0.1 – 1.0 µs first interacted with each other, forming TM dimers (Fig. 1). To follow the association of the TM peptides, we monitored the distance between the center of mass (COM) of the helix monomers for all the 11 receptors (Fig. S1). The TM peptides for all the 11 receptors dimerize in the DMPC bilayer and appeared stable throughout the simulation, in that few, if any large scale helix separations (> 20 Å) are seen for the remainder of the simulations.

**Figure 1.**
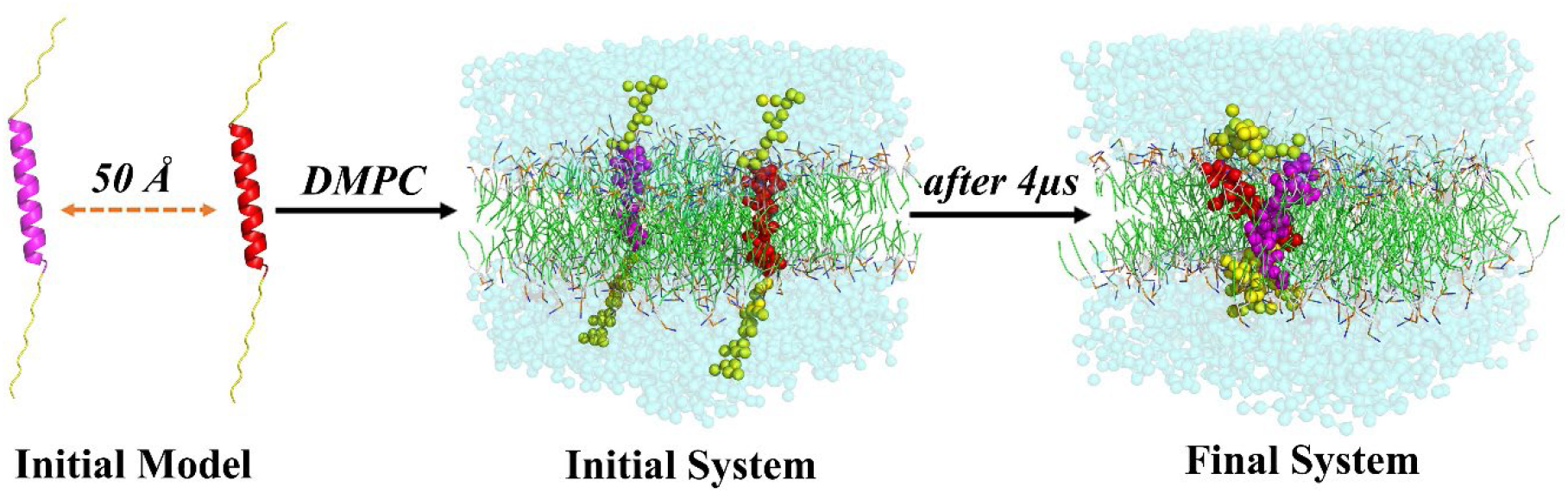
Association of TM dimers in the DMPC bilayer. TM peptides are initially placed 50 Å apart from each other and then inserted into DPMC bilayer. TM peptides interact quickly and the final 4 µs dimer conformation is shown here. TM region of the peptides are shown as magenta and red. N- and C-terminal regions are shown as yellow.

For Martini 2 it became known that the model is slightly too sticky for protein-protein interactions in the membrane^22–24,34^ and it is therefore important to provide a negative control. This is made difficult due to the absence of experimental data, to our knowledge for any of the 11 TM peptides studied. We examined polyLeucine (K_2_-L_26_-K_2_) as a general control as it prefers to stay in monomeric form in bilayer environment.^35,37^ Indeed, this peptide shows higher population of the monomeric state in the bilayer with Martini 3 compared to the previous Martini 2 model.^26^ Multiple simulations carried out for 4.0 μs, under the conditions of the study presented here, also showed no interaction between the polyleucine peptides using Martini 3 (Fig. S2). The polyLeucine peptide did interact, albeit qualitatively more weakly, with some of the 11 TM peptides tested in the simulations, suggesting that a generic peptide as a general negative control peptide may not be appropriate. The topic of negative controls, is a topic which has been largely avoided in the literature and progress in this arena warrants its own report.

To simplify the analysis of the systems, all 4 repeat simulations for each receptor TM dimer system were concatenated and total populations of the TM dimers are calculated considering the inter-helical angle and distance as shown in the 2D plot for Glycophorin A and EphA1 as well as EphA2 (Fig. 2A-C). Each 2D plot has one global minimum (highly dense population) and 2/3 local minima (2D plots for the other 9 systems are shown in Fig. S3). In few cases (ErbB1, ErbB1/B2, ErbB2), we saw the distribution of one large global minimum, with modest indication of side-minima, but nevertheless, we divided this into three groups in order to look at the other structures for possible alternate configurational states. We then selected up to 5 representative structures from the geometric center of each of these 3 clusters for each of the 11 TM proteins, comparing them with the NMR structure for further analysis. The best ones, in terms of crossing angle and root mean square deviation (RMSD), are reported in Figure 3, the average and standard deviations are shown in Table 1. Interestingly, using only the non-polar sequences of the TM helices did not yield stable dimers in many of the 11 systems (data not shown) using Martini 3 CG simulations, again pointing to the importance of the flanking regions. By contrast, the webserver PREDDIMER is a method which is able to reliably deal with TM domains only and produces model structures close to those derived by NMR for the 11 systems (Table S3)^6,19,20^ and reasonable results for other systems.^28^ This approach is based on the interhelix alignment of the peptide’s surfaces, with the possible dimers ranked by a parameter, Fscor, which considers the best packing and non-polar/hydrophobic complementarity between peptides and exposed dimer surface to lipids. Most important here is that PREDDIMER also gives several models (i.e. alternate configurations) which can be compared with the structures from the CG simulations (Table S3). As an additional comparison, we also include predictions using Alphafold2 Multimer for the sequences with N- and C-terminal extensions in Table S2, but considering only the models with the highest confidence values (Table S4).

**Table 1:** Comparison of CG simulated TM dimers with the NMR structures. Central conformers of the three most populated clusters from the CG simulation are considered for calculating the mean and SD of the crossing angle values. Similar/ near crossing angle (X) values (± 20°) and RMSD ≤ 3.5 Å between the CG and NMR are marked as red.

| TM Dimers | CG simulation |  |  |  |  |  | NMR |
| --- | --- | --- | --- | --- | --- | --- | --- |
|  | X (deg) |  |  | RMSD (Å) from NMR |  |  | X (deg) |
|  | 1<sup>st</sup><br>Cluster | 2<sup>nd</sup><br>Cluster | 3<sup>rd</sup><br>Cluster | 1<sup>st</sup><br>Cluster | 2<sup>nd</sup><br>Cluster | 3<sup>rd</sup><br>Cluster |  |
| 1-Bnip3 | -60 ± 10 | -87 ± 4 | -89 ± 5 | 4.1 ± 0.5 | 5.2 ± 0.3 | 5.4 ± 0.3 | -41 |
| 2-EphA1 | -47 ± 5 | 49 ± 6 | -15 ± 10 | 3.5 ± 0.2 | 6.5 ± 0.2 | 4.8 ± 0.1 | -52 |
| 3-EphA2 | -11 ± 3 | 29 ± 4 | 33 ± 8 | 5.1 ± 0.7 | 5.2 ± 0.5 | 4.6 ± 0.1 | 17 |
| 4-ErbB1 | -30 ± 2 | -8 ± 6 | -20 ± 8 | 3.3 ± 0.4 | 3.9 ± 0.1 | 4.3 ± 0.3 | -42 |
| 5-ErbB1/2 | -16 ± 4 | -55 ± 5 | -26 ± 5 | 4.0 ± 0.2 | 3.6 ± 0.3 | 2.4 ± 0.9 | -51 |
| 6-ErbB2 | -60 ± 4 | -18 ± 5 | -88 ± 8 | 2.7 ± 0.9 | 4.1 ± 0.1 | 3.6 ± 0.6 | -43 |
| 7-ErbB3 | 40 ± 11 | -8 ± 4 | -41 ± 9 | 3.9 ± 0.1 | 4.5 ± 0.2 | 5.1 ± 0.3 | 30 |
| 8-ErbB4 | -30 ± 4 | -56 ± 4 | -10 ± 6 | 3.9 ± 0.4 | 4.1 ± 0.2 | 5.4 ± 0.3 | -41 |
| 9-FGFR3 | -18 ± 8 | -50 ± 3 | 34 ± 5 | 5.6 ± 0.1 | 5.2 ± 0.2 | 5.6 ± 1.2 | 35 |
| 10-GpA | -19 ± 5 | 0 ± 4 | 14 ± 3 | 3.0 ± 0.2 | 4.6 ± 0.2 | 3.4 ± 0.3 | -39 |
| 11-PDGFRb | -84 ± 3 | -132 ± 7 | -61 ± 8 | 5.5 ± 0.4 | 6.9 ± 0.1 | 5.3 ± 0.5 | 23 |

**Figure 2.**
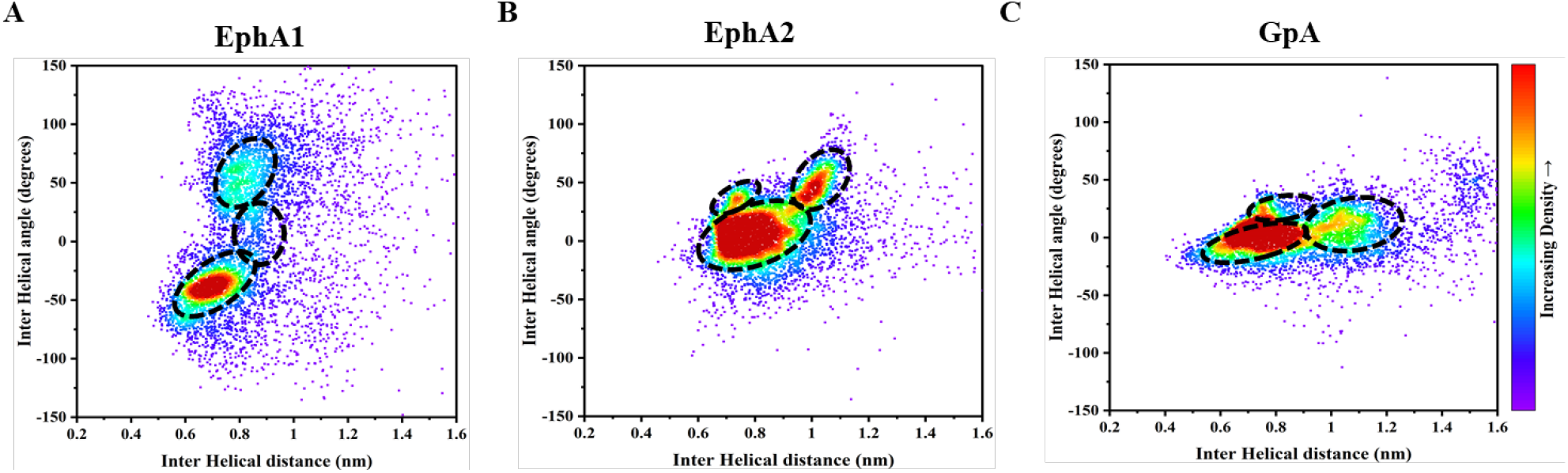
2D distribution plot (interhelix angle vs. distance) for (A) EphA1 (B) EphA2 and (C) GpA. Each plot is divided into 3 population clusters based on the inter-helical angle and the inter-helical distance, as described in Methods. The population of each cluster are shown in Table1. Several structures from each population cluster was extracted and then their average compared with the NMR structure (shown in Table1). Data from the last 2.5µs simulations are considered. Data points at intervals of < 500ps are skipped for clarity.

**Figure 3.**
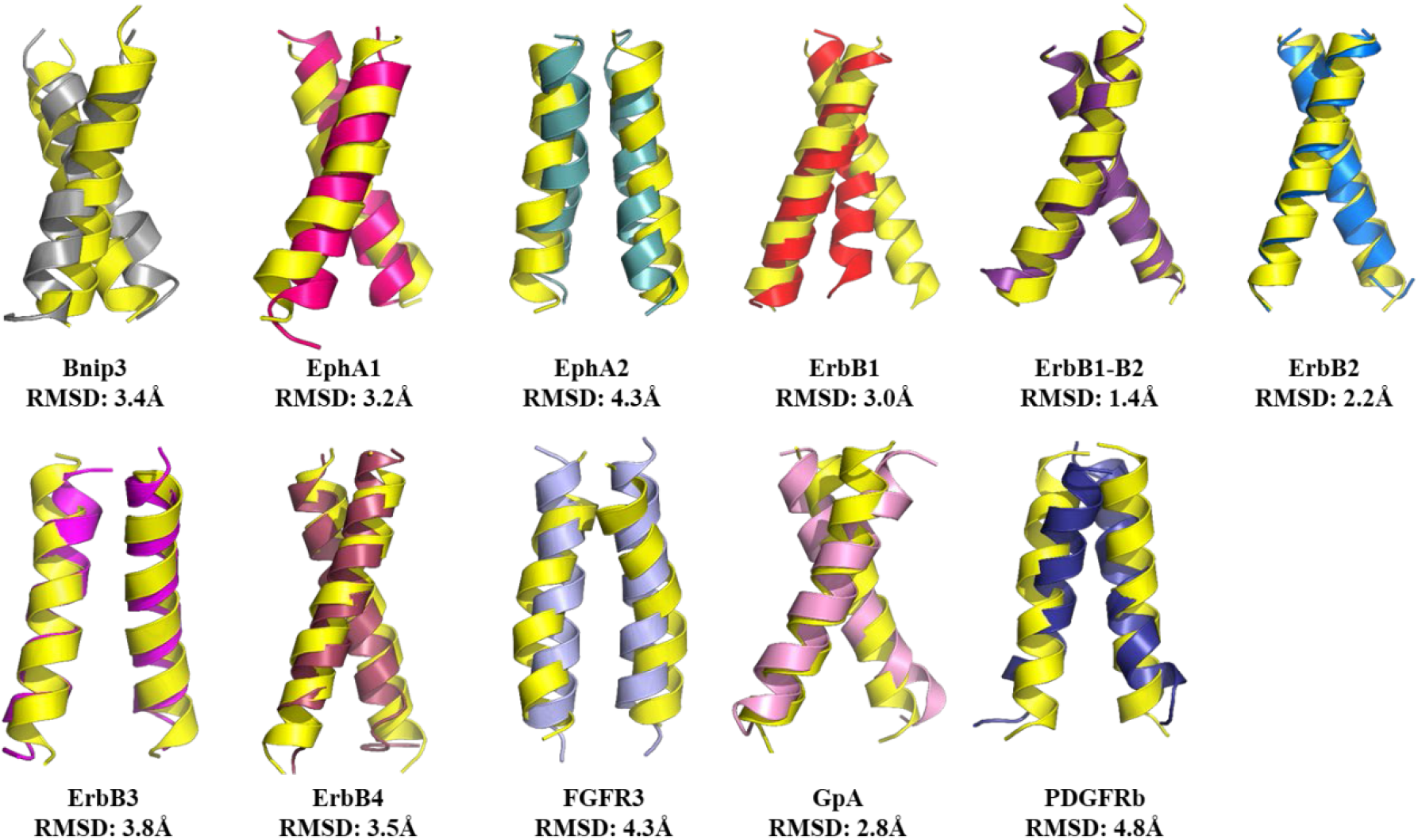
Superposition of the 11 solution NMR TM dimer structures (in yellow) with best fit CG simulated structures (vars. colors). All the CG structures are converted to all-atom (AA) representation. Best fit structure represents the one (among the 5 structures picked from each of the clusters) that has lowest backbone RMSD from the NMR structure. Only the non-polar helical TM region is considered for backbone RMSD calculation.

The comparison of the RMSD of the CG simulated TM dimers with the NMR structures (Fig. 3, Table 1), using the non-polar TM helical region of the structures only, shows that the CG structures range from nearly similar^36,38,39^ (2.4 Å for ErbB1/ B2 dimer) to more distant (5.3 Å for PDGFRb). The same is the case for the *ab initio* PREDDIMER models (Table S3) where the RMSD with the NMR structure also ranges from 1.7 Å (for GpA) to very distant 9 Å (for ErbB1/2). It should be noted that in general the CG structures compare less well with the NMR structures than the best PREDDIMER models, suggesting that the flanking region have an effect on this comparison for some of systems, as well as dynamics (see below). Specifically, EphA1 and –A2 as well as ErbB4, FGFR3, GpA and PDGFRb are all predicted well (< 2.5 Å backbone RMSD) by PREDDIMER whereas the CG simulations all yield structures with RMSDs > 3.5 Å for these, except 3.0 Å in case of GpA. Nevertheless, most of the CG simulated TM dimers (10 out of 11) share, within +/- 20°, similar values of crossing angle with that of the NMR structures (Table 1). The exception is PDGFRb, where the CG TM dimer configurations are right-handed (a negative value of the crossing angle) compared to left-handed (a positive value of the crossing angle) NMR dimer structures. It is also remarkable that not always the lower energy (best Fscor) or first PREDDIMER model corresponds to the NMR structure (as is the case for EphA2, ErbB1, ErbB1/B2 and PDGFRb, where the second model is the better one. This was observed in a previous PREDDIMER study as well,^19^ even though the PREDIMER method has been refined since.

As expected, given its recent success, Alphafold2 Multimer showed the best comparison with NMR structures, with lower RMSD values than those of PREDDIMER and especially Martini 3 structures (Table 2). In using Alphafold2, 9 of the 11 structures predicted are within 3.5 Å of the NMR structure (7 structures within 1.5 Å). By contrast 6 of 11 CG TM structures are similar to the NMR structures, with none of the cluster center structures less than 2.4 Å). Despite this good agreement for Alphafold2, however, some dimers (with RMSDs equal or lower than 1.0 Å to the NMR structures) show relatively low confidence scores, in the range of 0.3-0.4, for AlphaFold2 Multimer (for instance ErbB3, ErbB4 and FGFR3). Alphafold2 Multimer confidence scores of > 0.5 are thought to be reliable structures – on the other hand, EphA1 which has an RMSD of > 6Å has a score of 0.5. Indeed, one may think that Alphafold could be highly influenced by the TM helix dimer structures which exist in the PDB (of which the 11 examples here form a significant subset). In order to test the possibility that the neural network simply remembered facets of these structures, we also ran the program when it was trained on the PDB only having structures of 2001 and prior (Table S4). Since all TM dimer structures were determined after this date (even the first GPCR multi-TM membrane crossing structure was published only in 2001), this could have led to a different/less biased prediction. However, the prediction is very similar, in fact significantly better for some TM dimers (RMSDs for EphA1 and ErbB1, now 1.3 and 0.9Å compared to 6.6 and 4.2Å, respectively for the 2020 database) whereas some are worse (e.g. ErbB3 3.6Å vs. 1.0Å, but both at 0.4 confidence). This result, surprisingly suggests that most information needed to accurately predict TM helix dimer structures was already present in the PDB in 2001, although few membrane structures had been solved. This implies that the structures of non-polar helices in the interior of soluble proteins may not be that different from the helixes of TM crossing proteins, to a large extent obeying the rules of coiled-coil packing.^40^

**Table 2:** Comparison of Alphafold2 multimer predicted structures (database up to 2020) with NMR and CG simulated TM dimers. Best representative structures from the three populations of the CG simulation and the top predicted dimer conformations are compared. Similar/ near crossing angle (X) values (± 20°) and the RMSD ≤ 3.5 Å with NMR structures are shown as green. Similar RMSD values with the CG structures are shown in blue. Higher confidence score values (>0.5) are considered best predictions and are shown in red.

|  | Alphafold 2 Prediction (Database upto 2020) |  |  |  | NMR |
| --- | --- | --- | --- | --- | --- |
|  | X (deg)/ Fscore | RMSD (Å)<br>from NMR | RMSD (Å)<br>from CG | Confidence<br>score | X (deg) |
| 1-Bnip3 | -38/ 3.9 | 0.6 | 4.0/5.2/6.1 | 0.6 | -41 |
| 2-EphA1 | 25/ 2.4 | 6.6 | 5.7/3.1/4.7 | 0.5 | -52 |
| 3-EphA2 | 11/ 2.3 | 1.0 | 5.3/5.0/5.1 | 0.5 | 17 |
| 4-ErbB1 | 21/ 1.8 | 4.9 | 5.6/5.4/5.6 | 0.4 | -42 |
| 5-ErbB1/2 | -27/ 2.4 | 3.4 | 2.9/2.4/3.0 | 0.3 | -51 |
| 6-ErbB2 | -18/ 1.6 | 3.3 | 4.4/3.9/5.5 | 0.4 | -43 |
| 7-ErbB3 | 17/ 2.0 | 1.0 | 4.7/5.0/4.7 | 0.4 | 30 |
| 8-ErbB4 | -42/ 3.6 | 1.1 | 4.6/3.5/5.8 | 0.3 | -41 |
| 9-FGFR3 | 37/ 3.5 | 0.7 | 5.8/4.9/5.7 | 0.4 | 35 |
| 10-GpA | -37.6/ 3.5 | 0.6 | 3.1/4.2/3.5 | 0.6 | -39 |
| 11-PDGFRb | 18/ 2.2 | 0.4 | 5.6/7.2/5.0 | 0.6 | 23 |

How do relatively large RMSDs arise in the comparison of the non-polar helical segments of the TM regions? It is instructive to consider the meaning of RMSD when comparing TM helix dimers with different crossing angles and helix rotations. Fig. S4 gives a RMSD reference for changes of an ideal modeled parallel helix dimer of EphA1, illustrating that RMSD values quickly become considerable upon rotation of one or both helices and crossing angle changes, even though the correct segment of the helices (N- vs. C-term) may still be in contact. Models such as these did not allow us to pin-point a systematic difference between AlphaFold2 Multimer and PREDDIMER models and those generated by CG Martini 3 peptide association simulations (effects due to the Martini 3 particle vs. all-atom representation are considered removed upon conversion to the latter, used prior to all analyses). However, the difference could simply arise from wider conformational energy landscapes/basins sampled in the CG simulations.

Remarkably, the PREDDIMER models share similarities with both CG simulated and the NMR derived structures (Tables 1 and S3). As shown in Fig. S5, the best fit structure from the 3^rd^ cluster of PDGFRb from simulation is a right-handed dimer with a larger crossing angle of −55° and shares close similarity (RMSD 3.2 Å) with the 3^rd^ predicted model of PREDDIMER. By contrast the NMR structure for this protein’s TM domain is left-handed dimer with crossing angle of 23° which shares similarity (RMSD 2.3Å) with the 2^nd^ predicted model. As the *ab initio* PREDDIMER program^19^ predicts different possible arrangements between the TM peptides, our CG simulation in most cases also provides similar information represented by the best fit configurations from the three most populated clusters. (Tables 1 and S3). Specifically, PREDDIMER and less so the CG simulations produce alternate configurations for the majority of systems, often as side clusters of higher energy (lower Fscor) and population, respectively. Our mid. 2022 Alphafold2 Multimer version is not yet sampling nearby states/ensembles, but this may be available in the nearer future.^41^ For the simulations, a critical question is whether these are local minima in which the simulation became “stuck” or whether there is adequate sampling and convergence (i.e. transitions between configurational states allowing their equilibration).^42^

The concern that the CG simulations are insufficiently converged is lessened by the observation that nearly identical regions of the 2D interhelix distance – crossing angle plots are sampled in all of the 4 trajectories and that we observe a considerable number of transitions (∼ 10 or more) between states with different crossing angles. Representative examples are shown for EphA1, A2 and GpA in Fig. 4 (with the full set of plots in Fig. S6-S8). In most cases, we observed transitions between the clusters for all the receptors (Figs. 2 and S3). Moreover, the analysis of the overall TM crossing angle distribution (Fig 4 and Fig S6-S8) for all the 11 TM dimers shows the existence of both the right- and left-handed configurations. By contrast, the NMR structures of each TM dimer only provide one configuration, resolved at particular pH and in a suitable bicelle or micelle environment. Therefore, as shown in Table 1 and S3, the TM models obtained from the CG simulations go hand in hand with *ab initio* PREDDIMER predicted models and share similarity with the NMR structures in terms of RMSD, Fscor and inter-helical crossing angle (Figs. 3 and S5). However, there are also examples of less than perfect agreement in case of Bnip3, EphA2, FGFR3 and PDGFRb. Possible origins of these differences are discussed below.

**Figure 4.**
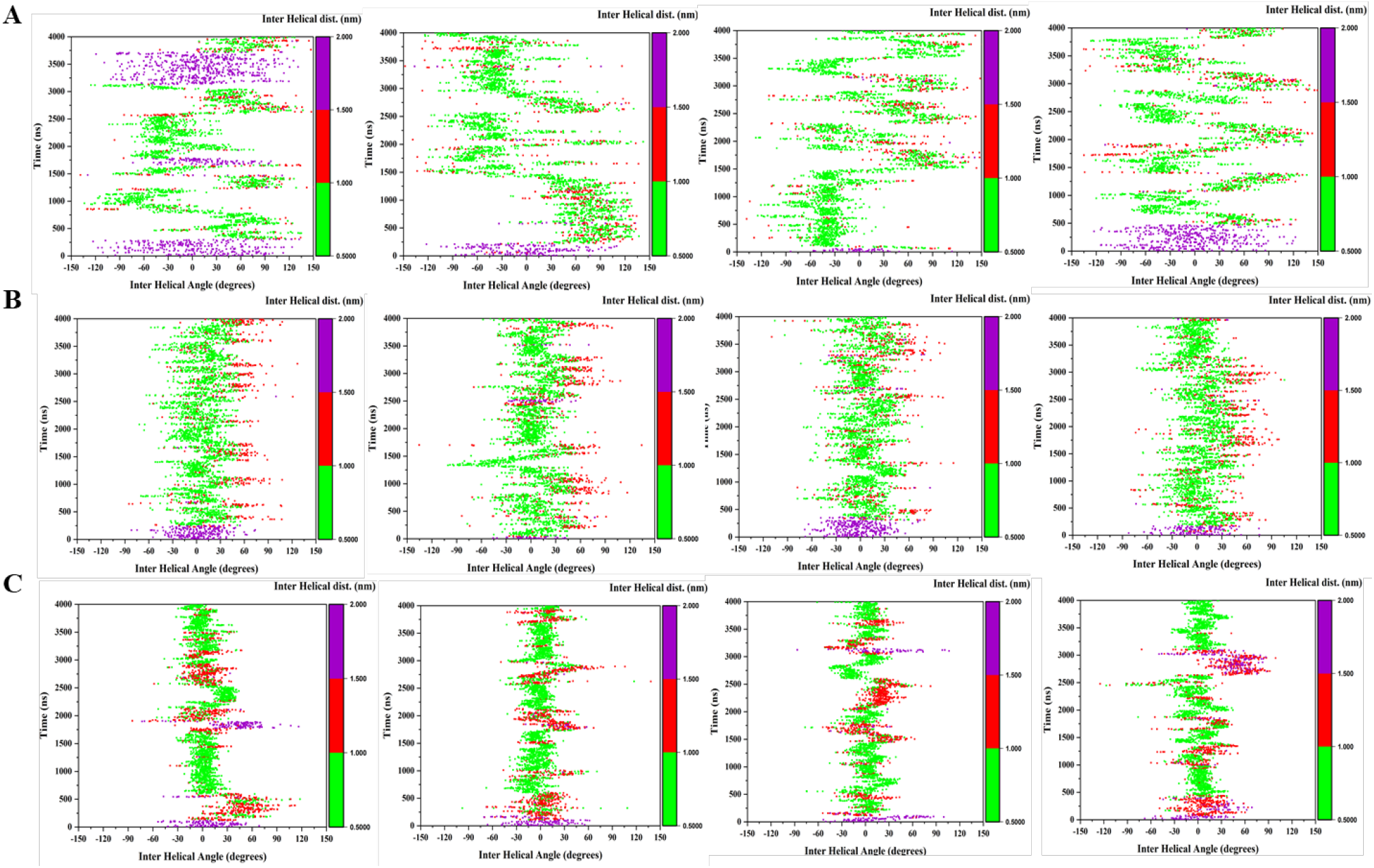
2D Plots showing the conformational transition of TM dimers over the simulation time considering the inter-helical angle vs inter-helical distance for (A) EphA1, (B) EphA2 and (C) GpA. Results from 4 trajectories are shown here. The plots are colored based on the inter-helical distance from 0.5- 1.0 nm (green), 1- 1.5 nm (red) and 1.5- 2.0 nm (purple).

A switching between the interaction interface of the TM regions results in different helix orientations in the membrane environment which needs to be coordinated with the proximity of the helices as it is typically difficult to have large changes in crossing angles without a temporary at least partial separation of helices (since larger sidechains would be blocking the transitions). To measure the change in TM orientation during the simulations, we calculated the 2D distribution of helix dimerization as a function of inter helical distance and the inter helical crossing angle (Figs. 2 and S3) and also plotted the contact map for the obtained TM dimers (Fig. S9). As expected, based on population density, both left-handed or right-handed configurations of the TM dimer are seen in some of the cases, most clearly for EphA1 (but several configurations-even though some are relatively close in crossing angle- are seen for most). The dimeric interface of the TM peptides from our study shows a high level of similarity with the NMR structures (Fig. S9). All the TM dimers except ErbB3, FGFR3 and PDGFRb contain a canonical extended GxxxG or SxxxG motif, starting from the middle of the sequence to the C-termini of the TM domains (Table S2). This motif allows for potential stabilization by Cα-H hydrogen bonding in the typical GAS_right_ dimer motif.^4,43^ In fact, as shown in Figs. 2, 4 and Figs. S3, S6-S8, EphA1, ErbB3 and ErbB4 showed an equal population of both right-handed and left-handed TM dimer configurations whereas Bnip3, ErbB1, ErbB1/ B2, ErbB2, FGFR3 and PDGFRb showed predominantly right-handed TM dimer configurations. Moreover, EphA2 and GpA demonstrated mostly parallel dimer configurations with a range of both right- and left-handed configurations. More information on the salient features of the CG derived structures is given based on simulation averaged and individual structure contact maps in the Supplemental Information (text, Fig. S9 and Table S5). Here we can comment on cases where information on alternate structures is available. Experimental data and extensive simulation data have been published on only a few of the systems: TM helix dimers of EphA1, EphA2, ErbB1. In all cases the TM helices dimerize in two different ways: either right-handed, left-handed helical dimers and/or with different areas of contact on the helices.^9,10,12–14^ Specifically, the NMR structure of EphA1 dimer is right-handed using an N-terminal ^550^AxxxGxxx^558^G motif at a low pH of 4.3, whereas there is partial unwinding in this region, and at pH 6.3 a more C-terminal GCN4-like motif is used involving ^560^Axxx^564^G.^9^ Whereas this represents essentially a 180° rotation of both helices the differences in contacts are more subtle, however, following molecular dynamics relaxation in the original paper. Extensive simulation work by the Sansom group using a locally modified version of the Martini 2 forcefield, reproduced many aspects of the experimental data, but had a second stable state which utilizes a ^559^AxxxLxxx^567^V motif.^25^ The second state 2 structure is more parallel compared to the NMR-like state 1 (N-terminally crossed) structure. In the CG simulations we observe contacts which encompass a mixture of both state 1 (on one helix) and state 2 (on the second helix). This asymmetry of the individual structures (in cluster 1, Table. S5) explains the rather modest RMSD of 3.5 Å with respect to the NMR structure. Cluster 3 has a lower right-handed crossing angle, whereas cluster 2 structures are unlike state 2 structures. They are asymmetric and have a left-handed crossing angle (Table 1).

The determination of the EphA2 NMR structure at pH of 5.0 revealed contacts via a left-handed configuration at a crossing angle of 15° and utilizes an extended heptad motif ^535^LxxxGxxAxxxVxx^549^L. However, a glycine zipper motif is also present in the TM sequence as ^536^AxxxGxxx^544^G which would be compatible with a −45° right handed structure, but is not seen experimentally.^10^ However, point mutations and kinase activity measured in cells with and without ephrin ligand had confirmed this model, switching between heptad and glycine zipper stabilized structures.^45^ Again, extensive simulations using the earlier Martini potential function recapitulated the close to parallel configuration (crossing at ∼ 10°) with the experimentally characterized heptad-like contacts,^47^ but the structures could also transition to a right handed configuration (with a crossing angle around −20°). This configuration could be stabilized in that report by mutating out parts of the heptad motif. Our cluster 2 and 3 structures have left-handed crossing angles but they are larger than those of the NMR structure (29-33° vs. 17°). Cluster 1 has structures which are more parallel with a slight right-handed crossing (−11°). None of the structures picked from the clusters are close to the NMR structure but the main-chain sites in closest contact involve the N-terminal ^540^Gxxx^544^G. For another project we carried out EphA2 TM dimerization simulations with Martini 3 but in a POPC bilayer and at pH 4.5, and found that a right-handed structure with the glycine zipper motif was predominantly populated. This was also the structure which was stabilized by TM-like peptides which were found to activate the receptor in a ligand independent manner by adding the peptide helix to the side of the TM dimer.^58^ We can at this point only speculate-but the left handed/parallel structure appears to be stabilized by Phe-Phe interactions which seem to be recognized by PREDDIMER and especially Alphafold2 (giving a 2.0 Å and 1.0 Å RMSD model, respectively to the NMR structure).

The EGFR TM has also been subject to extensive experimental and simulation studies. The first NMR structure determined in DPC used a C-terminal ^637^Axxx^641^G motif and is thought to be inactive, while the N-terminally crossed ^625^Gxxx^629^A motif, later determined in bicelle represents the active state^11,50^ and is used for comparisons here. Again, extensive metadynamics simulations with a version of Martini 2 were able to sample configurations close to these NMR structures. An energy landscape was mapped, showing that both N- and C-terminally crossed structures have similar right handed crossing angles and have facile pathways of interconversion by change of contact region (which can be accomplished without rotation of the helices).^24^ In the main cluster our Martini 3 simulations only came within 3.3 Å on average to the NMR structure. Looking at the contacts of individual structures it is evident that considerable sliding must occur as, similar to EphA1, the N-terminal contacts and C-terminal-like Glycine-zipper residues are populated on the two helices separately, meaning that the most stable contacts are made in the middle of the TM dimer, as is evident from the trajectory average contact map (Fig. S9). In fact, the N-terminal GxxxG motif seems to continue into a GCN4-like motif on one of the helices, while the other helix has slid relative to the first one and employs a GCN4-like motif towards its C-terminus (where a Phe – Gly interaction between the helices may play a role). When larger juxtamembrane regions are included, this may help to stabilize particular structures from such mixtures – specifically the N-terminal part of the JM region is known to form an antiparallel short helix dimer as shown experimentally and by NMR.^29^ It should be noted that in a POPC membrane the RMSD of cluster 1 structures in a Martini 3.0 CG simulation of a similar EGFR TM region is 1.7 Å with similar crossing angle (right handed 27°) and that the asymmetry effects are lessened.^52^ As would be expected, the CG structures which compare best with the NMR structures (ErbB1/B2 heterodimer and ErbB2 homodimer) are more symmetric also. We may speculate that ErbB1/B2 and ErbB2 dimer, as well as ErbB3 and –B4 could also form C-terminally crossed inactive structures, but these remain to be studied/reported. We skip the example of GpA (again a reasonable prediction at 3.0 Å RMSD) as a functional role for any alternate structures is as yet unknown.

As a further examination of the different dimer binding modes, we calculated the free energy of binding for EphA1, EphA2, ErbB2 and for the well-studied helix dimer of GpA^42,44,46,48,49,54^ starting umbrella sampling (US) runs from each of the three cluster centres as well as from the NMR structure. We found that the potential of mean force derived free energy estimates are similar despite using presumably higher energy starting structures (for details, see Figs. S10 and S11 and additional discussion in the supporting material). Several studies also suggested that the lipid environment and also presence of cholesterol may control the mode of dimerization.^17,20,32,33,51^ For example, it is known that negatively charged lipids, such as POPS and cell signalling lipids PIP2 and PIP3 alter the functional behavior of TM-but also TM peripheral proteins cells.^53,55,56^ Specifically, it is noticeable that almost all of the 11 systems chosen here contain a cationic “plug” to prevent sliding of the C-terminal region into the membrane and recent work by Barerra and colleagues^31^ suggests that EphA2 may switch from a parallel TM helix dimer in a wider membrane with PIP2, where positive charges can be tolerated to the structure with a wider crossing angle in a thinner membrane without PIP2, where the juxtamembrane regions may repulse. In this sense the NMR studies of and the predictions/simulations of TM dimers here seem rather artificial since only neutral zwitterionic detergents/lipids were used.

The overall picture which emerges from the CG simulations in this study (but is supported by PREDDIMER *ab initio* models) is that TM helix dimers of most of the 11 systems examined assume a preferred configuration but that side-minima and in many cases where NMR derived structures are available, alternate configurations of the dimer are also sampled (see above). Importantly, these structures can interconvert on the μs-CG-timescale without requiring a complete separation and rebinding of the TM helices. This suggests that TM dimers by themselves are not as locked into one configurational state as one might have perceived from NMR studies, which are –by necessity-presenting a relatively tight ensemble of a single conformer. Alphafold2 Multimer predicts well defined NMR-like structures given the AI nature of approach, which can often reproduce the experimental structures present in the PDB dataset at crystallographic resolution (< 1.5 Å). Which particular conformer is stabilized or destabilized by the further addition of TM domain flanking residues and especially charged lipids needs further study, as does the integration between TM domain configurations and the protein-protein as well as protein-membrane interactions of lipid-bilayer proximal extracellular and intracellular domains. Still, the experimental results on the efficacy of TM-like activating and inhibiting peptides on some of the type-1 receptor systems^52,57,58^ as well as key TM localized cancer mutants^29,59^ suggest that configurational equilibria involving the TM region has a considerable influence on the function of the TM proteins.

Caution needs to be exercised when comparing *ab initio* predictions, MD simulation results or even structures derived from experiments. First, the comparison needs to be performed for the same TM peptide or protein construct, which is not the case here. Also, we did not see value in doing calculations at unphysiological low pH (Table S1 and S2). Second, membrane model system composition is important and some of the NMR structures were determined in DPC detergent, rather than in bicelles, but there is no systematic difference among this set of examples. However, even bicelles are not the idealized DMPC-composed planar bilayer in the middle, with DHPC detergent at the bicelle-disk edge, as some studies are showing a peptide dependent mixing of the DHPC and DMPC molecules. Third, the NMR derived structures are biased by restraints between the two helices which are for the most part symmetric. By contrast, in the simulations a sliding of helices relative to one another is observed (most noticeably for EphA2 and ErbB2), especially when individual structures are examined. It is worth remarking that *ab initio* predictions as PREDDIMER (and Alphafold2 Multimer) cannot consider variations in membrane thickness, lipid composition or salt concentration in the aqueous solution, while MD simulations with Martini 3 can naturally include these environmental effects. Even differences in pH can be mimicked by defining different charged states for acidic/basic groups, or by using the Titratable Martini approach.^60^ Moreover, trimers or oligomer can be accurately studied^58^ with Martini 3, the model is adapted to simulate bilayers of different composition^67^ or interactions in larger protein complexes.^61^ While Wade and colleagues scale protein-water interactions down by 10%, this does not lead to better defined structures for GpA (or for TrkA) suggesting that such a blanket approach may not solve Martini 3’s remaining limitations. Additional simulations and experimental studies are needed to delineate the possible effects of such variables on the TM dimer structural configurations and their stability.

In conclusion, we have shown with an established set of type-1 TM proteins of interest to the structural community, that Martini 3 CG simulations can predict the structure of TM dimers, as well as their tendency to form alternate structures, reflecting different specific residue-residue (motif) interactions, between the TM peptides. Our results suggest that TM domain flanking sequences are likely responsible at least for a shift in the population of TM configurational states which are sampled and that such sequences should be considered in the design of TM-like peptides which may inhibit or activate the TM protein’s function.

## Acknowledgements

This work is supported by a NIH R01 grant from the National Eye Institute R01EY029169 and previous grants from NIGMS (R01GM073071 and R01GM092851) to the Buck lab. We acknowledge Gilberto Pereira for supporting us with the AlphaFold2 Multimer predictions.

## Author Contributions

ARS generated the PREDDIMER predictions, the protein coarse-grained models and performed the MD simulations and analysed the data. ZM carried out early simulations with the beta release of Martini 3. PCTS contributed with critical data, discussions and guidance for the coarse-grained model generation and for analysis performed. MB conceived, supervised and led the project. ARS and MB co-wrote the paper, with contributions from PCTS. All authors read and approved the final version of the manuscript and supporting information.

## STAR Methods

### Data Sets and Modeling of the TM peptides

The NMR structures of 11 TM dimers were extracted from the PDB database, using the first structure of the ensemble. All the TM dimer structures were determined either in DPC micelles or DMPC/DHPC bicelles with the pH value ranges from 4.5 to 6.8 (Table S1). The TM region sequences for all the 11 proteins were extracted from the UniProt database and the UniProt definition of the TM hydrophobic domain was used (sequence in bold). The TM domain of the peptides was modeled as an ideal α-helix using PyMOL 2.4. We then added extra 8-9 residues at N-terminal and 10 residues at the C-terminal of the TM models as an extended conformation (Table S2) as these extra- and intracellular membrane proximal regions are known to provide better stability for some systems in the lipid bilayer.

### Coarse-grain molecular dynamics simulation

In order to characterize the dimerization of TMs, we built 11 TM peptide systems with the monomers placed 50 Å apart from each other (Fig. 1). For this, atomistic (AT) modeled systems of all the 11 TMs were converted to coarse-grained (CG) representation using the *martinize2*.*py* workflow module of the MARTINI 3 force field^26^ (see https://github.com/marrink-lab/vermouth-martinize) considering the secondary structure DSSP assignment.^62^ We used the elastic network to reinforce the stability of the helical secondary structure of the TM monomers. We used default values of the force constant of 500 kJ/mol/nm^2^ with the lower and upper elastic bond cut-off to 0.5 and 0.9 nm respectively. CG simulations were performed using GROMACS version 2016.5.^63^ The *insane*.*py* script^64^ was used for setting up of the DMPC bilayer (typically 306 lipids and 4870 CG water molecules) around the peptides in a cubic box with dimensions of 100×100×100 Å^3^. The pH of the systems was considered neutral. All the simulations were run in presence of regular MARTINI water and neutralised and brought up to 0.15M NaCl. The systems were equilibrated for 500 ps. The long-range electrostatic interactions were used with a reaction type field having a cutoff value of 11 Å.^65^ We used potential-shift-verlet for the Lennard-Jones interactions with a value of 11 Å for the cutoff scheme and the V-rescale thermostat with a reference temperature of 320 K in combination with a Berendsen barostat with a coupling constant of 1.0 ps, compressibility of 3.0 × 10^−4^ bar ^-1^, and a reference pressure of 1 bar was used. The integration time step was 20 fs. All the simulations were run in quadruplicate for 4 µs. For further analysis and comparison, the extracted CG structures were then converted to all atomistic (AA) representation using the Backward tool^66,70^ of Martini (as shown in Fig. S12).

### Data Analysis

The PREDDIMER webserver was used both to predict models for the 11 TM dimers *ab initio*, but also to analyse the NMR and CG MD derived structures based on the Fscor and helix crossing angle.^19^ Alphafold2 Multimer^30^ was used to predict dimer models for the same set of complexes, using a local installation of the ParaFold pipeline.^68^ This installation was recently validated by Martin and colleagues in 2022,^69^ reproducing the results predicted by Evans et al., 2021.^30^ Only the best model with the highest confidence value was used for the comparisons here. Analysis of the trajectories was carried out using tools in GROMACS.^63^ All the analyses have been carried out considering only the non-polar (uniprot defined) TM region of the peptides. The contact maps between the helices (again non-polar TM regions only) were calculated with a distance cut off 6 Å for all the backbone atoms. Interhelix distances are calculated between the center of masses of the TM, but not of the N- and C-terminal regions. 2D plots were plotted in Origin2020b using the inter-helical distance and the inter-helical angle between the TM helices, as x- and y-axes respectively. Each plot shows the presence of one global minimum and up to 2 local minima and we therefore divided into three populations in all cases. except for ErbB1, ErbB1/B2 and ErbB2. In these three cases, we saw the distribution of one large global minimum, with modest indication of side-minima and therefore decided to divide the global minimum into three groups in order to look at other near-by structures for more sampling. In all cases, for each of the three populations/groups, 5 representative configurations were extracted near the population maxima and then further compared with the experimentally derived NMR structures. The best one, in terms of RMSD and crossing angles, was then reported for the 3 minima (Table 1 and Fig. 3). The comparison with the NMR structure is done by calculation of backbone RMSD considering only the non-polar helical TM regions of the receptors (which varies less than 1.5Å within the NMR ensembles), whereas the N- and C-terminal additional residues are more flexible. We have also added the initial parameters for the 11 systems in the GitHub repositories for reproducibility (https://github.com/amita-bucklab/TM-dimerization).

## Supporting Information Available

Additional results, methods, supporting tables and figures are included in the supporting information

