## Supplemental info for "Transmembrane region dimer structures of Type 1 receptors readily sample alternate configurations: MD simulations using the Martini 3 coarse grained model compared to AlphaFold2 Multimer"

### Materials and Methods:

#### *Calculation of Interhelical Crossing Angle*

For the calculation of the interhelical crossing angle between the TMs, we used the method described in this paper.[1] We considered two points in each helix X and Y. X1 and Y1 are the center of mass (COM) of the helix X and Y (considered the Cα atoms) whereas X2 and Y2 are the COM of the top half of the helix X and Y respectively. The distance between X1 and Y1 is considered as the interhelical distance (d). Therefore, the interhelical angle (θ) is calculated as the dihedral angle between X2-X1-Y1-Y2. (Negative value of θ indicates right-handed dimer configuration and vice versa).

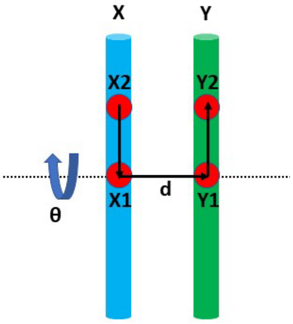


#### *Umbrella Sampling (US) simulations*

We used the umbrella sampling method with the center of mass (COM) distance between the TM peptides as the reaction coordinate and calculated the free energy of TM association of these receptor's TM domains. We calculated the dimerization free energy profiles for the TM peptides of GpA, EphA1, EphA2 and ErbB2, as examples using a standard umbrella sampling (US) protocol. The best fit structures obtained from the three most populated clusters (according to 2D plot in Figs. 2 and S3D) from the CG simulation were chosen as the starting point for the US simulations. Unlike the CG simulation above, the peptide length was reduced by removing the N- and C-terminal residues and we have considered only the transmembrane regions for all the US simulations. The TM dimers were embedded in the lipid bilayer consisting of typically 328 DMPC molecules, 5200 water beads with a salt concentration of ~0.15 NaCl using the same *insane.py*.[2] The center of mass distance between the TM domains was used as the reaction coordinate (ξ). US simulations were set up with 19 replicas spaced between the reaction coordinate of 0 to 3.6 nm with a spacing of 0.2 nm. The spring constant of 1000 kJ $mol^{-1}$ $nm^{-2}$ was employed for the TM domains. Each US window was simulated for 0.5 µs with a time step of 20 fs. The gmx wham[3] tool was used for calculating the potential mean force (PMF) profiles, estimating the free energy of TM domain dimerization. All the simulations were performed in GROMACS 2016.5. In order to calculate an average free energy and standard deviation between the US simulations begun with the three different conformers, the lowest energy minima values for each were weighted by the CG populations, derived from the three population clusters (listed in the legend of the 2D plot shown in Fig. S3).
The free energy of the TM dimerization ($\Delta_{ass}G_0$) was calculated from PMF[10] according to the following equations.

$$\Delta_{ass}G^0 = -RT ln K_{ass}$$

$$K_{ass} = \frac{\int_0^{r_c} 2\pi r e^{-\frac{PMF(r)}{RT}} dr}{A^0}$$

where $K_{ass}$ denoting the association constant, $A^0$ the standard area, and $r_c$ a cut-off.

**Additional observations for each of the TM systems:**

**Bnip3 TM dimerization:**

For Bnip3, the dimerization interface includes mostly the $^{168}$SxxxSxxxA$^{176}$ motif (Fig. S9A) and is similar to the NMR dimer where there is an additional stabilizing interface consisting of $^{180}$GxxxG$^{184}$ motif towards the TM C-terminus. Also, the comparison of the crossing angle (Figs. S3A and S6A) suggests that most of the dimer populations are in right-handed conformation.

**EphA1 & A2 TM dimerization:**

The CG EphA1 dimer is stabilized by both $^{554}$GxxxG$^{558}$ motif (Fig. S9B) similar to the NMR dimer and shows the presence of both the right-handed and left-handed TM dimer populations (Figs. 2A and 4A). However, all CG 15 structures looked at more closely in Table S9 are asymmetric. By contrast, EphA2 TM dimerization involves mostly symmetric contacts, but surprisingly these are not as well populated (defined) as in the case of EphA1 (Fig. S9C). The N-terminal $^{540}$GxxxG$^{544}$ motif is shifted relative to the $^{539}$GxxAxxxVxxL$^{550}$ zipper motif seen by NMR. and the hydrophobic patch of $^{546}$VxxLVL$^{551}$ as shown in Fig. S9C, which is also seen in the NMR dimer. Though the NMR dimer of EphA2 has a left-handed conformation, we saw both the left and right-handed conformations in the 2D plot (Figs. 2B and 4B) as seen in case of EphA1.

**ErbBs TM dimerization:**

On the other hand, ErbB1 (with a RMSD of 3.3 Å) has $^{649}$GxxxAxxxL$^{657}$ motif similar to NMR dimer (N-term. contacts) on one of the helixes but this is mixed with a C-terminal motif on the other helix (Fig. S9D) yielding asymmetric structures, as noted in the main text. The ErbB1/2 heterodimer (2.4-3.6 Å) shows the association with mostly a $^{649}$GxxGAxxxL$^{657}$ motif from ErbB1 and $^{656}$SxxxGxxxV$^{664}$ motif from ErbB2 similar to the interactions seen in the NMR structure. (Fig. S9E). 2D plot for the ErbB1/2 heterodimer shows the abundance of right-handed TM dimer conformations (Figs. S3C and S6C) similar to the NMR dimer based on the crossing angle calculations. In case of ErbB2 (RMSD of 2.7 Å), the TM homodimer self-associate through $^{656}$SxxxGxxxV$^{664}$ motif which is symmetric for cluster 1 structures (Fig. S9F) with the dimer population of mostly right-handed conformations similar to NMR conformer (Fig. S3D and S7A). As shown in Fig. S9G, the ErbB3 homodimer (RMSD 3.9 Å) associate mostly through a contact region which seems to have many variants, with only T47 standing out as well populated differing from contacts seen in the NMR structure. Intriguingly, the crossing angle distribution confirms the presence of left-handed dimer populated conformations similar to the NMR structure (Fig. S3E and S7B). With the backbone RMSD of 3.8 Å from the NMR structure, the dimerization of ErbB4 is predominantly right-handed but is asymmetric with one helix (chain B) utilizing a GxxGG motif (Figs. S9H, S3F and S7C).

**FGFR3 & GpA TM dimerization:**

The FGFR3 TM dimer is stabilized by well populated GxxxG-like and van der Waals and π-π stacking contacts involving $^{378}$SYxxGFFxF$^{386}$ similar to the NMR structure but shows mostly right and fewer left-handed preference by contrast to the left-handed conformations of the NMR structure (Figs. S9I, S3G and S8A), thus accounting for the high RMSD values (> 5.0 Å). The most thoroughly studied GpA dimer shows a right-handed association involving $^{79}$GxxxG$^{83}$ motif at its core similar to the NMR structure (Figs. S9J, 2C and 4C) with an RMSD of 3.0; a left-handed structure is also seen utilizing similar contacts (RMSD 3.4 Å).

**PDGFRb TM dimerization:**

In case of the PDGFRb homodimer, we saw predominant presence of right-handed conformers throughout the simulation and the association involves a canonical heptad repeat $^{536}$SAxxAxxVLxxI$^{547}$ similar to the left-handed NMR dimer (Figs. S9K, S3H and S8B), but the crossing angle difference seems to account for the large RMSD differences of > 5.0 Å. The CG structures are crossed significantly more at times (up to 132 degrees) for PDGFRb as the helices can insert up to 31 residues into the membrane, compared to GpA, where this number is more typical of TM regions with 23 residues.

**Free energy of TM binding:**

To get better insights into the stability of dimer binding modes, we calculated the free energy of binding for EphA1, EphA2, ErbB2 and for the well-studied helix dimer of GpA. The first structure (typically the one with the lowest energy or number of violations of experimental restraints) of the pdb-deposited NMR structural ensemble was taken as the starting configuration for the umbrella sampling. The other three starting structures represent the best fit structures extracted from the three population clusters, each cluster center (Table 1). We calculated the PMF along the reaction coordinate (COM between the TMs) for all the four TM homodimers (Fig. S10). Although the PMF calculations are computationally expensive compared to other methods [4] and we cannot be sure whether the simulations are well converged in all cases, the PMF technique has become a standard and, in this case, estimates free energy values, which are overall in good agreement with the experimental studies. In all the four cases, three PMF values show good agreement in terms of the lowest minimum PMF with slight difference considering the distance between the center of mass of the TMs which might be because of different initial configurations (Fig. S10). However, the unrestrained simulation shows extensive interconversion between cluster centers (e.g. Figs. 4 and S6-S8) and thus such interconversion is also likely in the even more extensive US calculations. The reported dimerization free energy of EphA1 is -15.4 ± 0.5 kJ/mol using the FRET studies.[5] Similarly, for GpA, the dimerization energy differs in a different membrane environment and it ranges from -15 kJ/mol to -31.5 kJ/mol.[6–8] As shown in Fig. S10, the average PMF values for EphA1 and GpA from our study are -9.2 ± 0.2 kJ/mol and -10.4 ± 0.2 kJ/mol respectively, which are reasonably close to the experimental values. Using the NMR structure as the starting configuration, the resulting PMF for the lowest energy interhelix distance of EphA1 and GpA corresponds to -8.2 kJ/mol and -13.5 kJ/mol respectively, which are comparable to the minima obtained from the three central structures of the CG simulation (Fig. S10A and D). For EphA2 and ErbB2, we obtained average PMF values of -11.1 ± 0.4 kJ/mol and -16.0 ± 0.3 kJ/mol respectively (Fig. S10B, C) considering the CG simulated structures which are similar (within a factor of 2 for EphA2 but much closer for ErbB2) with the PMF values obtained from the corresponding NMR structures (-6.3 kJ/mol and -17.6 kJ/mol respectively). The free energy of binding of all the four dimers are shown in Fig S11. Though the binding energy of full-length unliganded EphA2 has been estimated (-5 ± 0.2 kcal/mol)[9] before, the binding free energy of ErbB2 is unknown. A recent study also shows a high correspondence of dimerization free energy between the experimental values and the values obtained using the new Martini 3, contrasting with the old Martini 2 version that showed very high binding energy due to excessive association.[10–12] On the other hand, scaling the protein-lipid interaction has be done in the Martini 2 model, which improved binding energies and destabilized the protein aggregates, while preserving the membrane properties.[11,12]

**References cites in the SI:**

1. Li, P.-C., Miyashita, N., Im, W., Ishido, S., and Sugita, Y. (2014). Multidimensional

umbrella sampling and replica-exchange molecular dynamics simulations for structure prediction of transmembrane helix dimers. J. Comput. Chem. *35*, 300–308. 10.1002/jcc.23494.

11. Javanainen, M., Martinez-Seara, H., and Vattulainen, I. (2017). Excessive aggregation of membrane proteins in the Martini model. PLoS One *12*, e0187936. 10.1371/journal.pone.0187936.

12. Majumder, A., and Straub, J.E. (2021). Addressing the Excessive Aggregation of Membrane Proteins in the MARTINI Model. J. Chem. Theory Comput. 10.1021/acs.jctc.0c01253.

Supplemental Tables S1 to S5 and Figures S1 to S12 are provided.

**Table S1:** List of the 11 TM dimers studied. Experimental conditions for these TM dimers used in various NMR studies are also listed.

| Receptor (PDB ID) | Lipid Bicelle | pH |
|---|---|---|
| Bnip3 (2J5D) | DMPC/DHPC | 5 |
| EphA1 (2K1L) | DMPC/DHPC | 6.3 |
| EphA2 (2K9Y) | DMPC/DHPC | 5 |
| ErbB1 (5LV6) | DMPC/DHPC | 5.8 |
| ErbB1/2 (2KS1) | DMPC/DHPC | 4.5 |
| ErbB2 (2JWA) | DMPC/DHPC | 5 |
| ErbB3 (2L9U) | DPC | 5 |
| ErbB4 (2L2T) | DMPC/DHPC | 5 |
| FGFR3 (2LZL) | DPC | 5.7 |
| GpA (1AFO) | DPC | 6 |
| PDGFRb (2L6W) | DPC | 6.8 |

**Table S2.** TM peptide sequences for all the 11 TM domains used in this study. The central TM (membrane embedded hydrophobic) regions (in bold) used for this study are based on the UniProt database. TM sequence limits based on the NMR structures are underlined. All the NMR TM dimer structures considered for comparison in this study were either N- or C-terminal tagged.

| Protein (UniProt ID) | Peptide Sequence |
|---|---|
| Bnip3 (Q12983) | IFSAEFLK**VFLPSLLLSHLLAIGLGIYIG**RRLTTSTSTF |
| EphA1 (P21709) | SRGLTGGE**IVAVIFGLLLGAALLLGILVF**RSRRAQRQRQ |
| EphA2 (P29317) | EGSGNLAV**IGGVAVGVVLLLVLAGVGFFI**HRRRKNQRAR |
| ErbB1 (P00533) | TNGPKIPS**IATGMVGALLLLLVVALGIGLFM**RRRHIVRKRT |
| ErbB1/2 (P00533/ P04626) | CPTNGPKIPS**IATGMVGALLLLLVVALGIGLFM**RRRHIVRKR<br>CPAEQRASPLT**SIISAVVGILLVVVLGVVFGILI**KRRQQKIRKY |
| ErbB2 (P04626) | CPAEQRASPLT**SIISAVVGILLVVVLGVVFGILI**KRRQQKIRKY |
| ErbB3 (P21860) | LVLIGKTHLT**MALTVIAGLVVIFMMLGGTFL**YWRGRRIQNK |
| ErbB4 (Q15303) | TLPQHARTP**LIAAGVIGGLFILVIVGLTFAVYV**RRKSIKKKRA |
| FGFR3 (P22607) | EELVEADEAGSVYAG**ILSYGVGFFLFILVVAAVTLC**RLRSPPKKGL |
| GpA (P02724) | AHHFSEPE**ITLIIFGVMAGVIGTILLISYGI**RRLIKKSPSD |
| PDGFRb (P09619) | VPHSLPFKV**VVISAILALVVLTIISLIILI**MLWQKKPRYE |

**Table S3:** Comparison of *ab initio* predicted PREDDIMER structures with NMR and CG simulated TM dimers. Best representative structures from the three populations of the CG simulation and the top 3 predicted dimer conformations are considered for calculation. Similar/ near crossing angle (X) values (± 20°) and the RMSD ≤ 3.5 Å with the NMR structures are shown as green. Additional/alternative structures with crossing angle and RMSD values similar to that of CG simulated structures (values listed in Table 1) are shown in blue.

| | PREDDIMER | | | NMR |
|---|---|---|---|---|
| | X (deg) | RMSD (Å) from NMR | RMSD (Å) from CG | X (deg) |
| 1-Bnip3 | -50/ 50/ 45 | 2.9/ 6.0/ 6.0 | 2.4/ 8.3/ 5.8 | -41 |
| 2-EphA1 | -40/ -6/ 40 | 2.0/ 4.2/ 7.5 | 3.6/ 3.9/ 5.8 | -52 |
| 3-EphA2 | -20/ 30/ -60 | 5.3/ 2.4/ 6.2 | 3.3/ 6.0/ 4.9 | 17 |
| 4-ErbB1 | 6/ -15/ 40 | 5.3/ 3.0/ 6.7 | 4.5/ 2.9/ 6.3 | -43 |
| 5-ErbB1/2 | 30/ -35/ 60 | 7.0/ 3.0/ 9.0 | 6.2/ 2.3/ 8.5 | -51 |
| 6-ErbB2 | -60/ 60/ -25 | 4.5/ 8.9/ 2.9 | 3.7/ 7.5/ 2.5 | -43 |
| 7-ErbB3 | 15/ 45/ -25 | 4.6/ 4.8/ 6.2 | 3.4/ 4.3/ 3.5 | 30 |
| 8-ErbB4 | -35/ 25/ 55 | 1.7/ 5.9/ 7.0 | 3.8/ 5.7/ 3.9 | -41 |
| 9-FGFR3 | 30/ -15/ 35 | 4.3/ 5.8/ 1.7 | 5.7/ 3.3/ 4.4 | 35 |
| 10-GpA | -55/ -25/ 50 | 2.2/ 1.7/ 7.1 | 1.5/ 3.8/ 6.4 | -39 |
| 11-PDGFRb | 60/ 30/ -55 | 4.7/ 2.3/ 6.0 | 8.0/ 8.0/ 3.2 | 23 |

**Table S4:** Comparison of Alphafold2 multimer predicted structures (database up to 2001) with NMR and CG simulated TM dimers. Best representative structures from the three populations of the CG simulation and the top predicted dimer conformations are considered for calculation. Similar/ near crossing angle (X) values (± 20°) and the RMSD ≤ 3.5 Å with NMR structures are shown as green. Similar RMSD values with the CG structures are shown in blue. Higher confidence score values (>0.5) are considered best predictions and are shown in red.

| | Alphafold 2 Prediction (Database upto 2001) | | | | | NMR |
|---|---|---|---|---|---|---|
| | X (deg)/Fscor | RMSD (Å) from NMR | RMSD (Å) from CG | Confidence score | RMSD (Å) from AF (2020 database) | X (deg) |
| 1-Bnip3 | -34/4.0 | 0.6 | 4.0/5.2/6.1 | 0.6 | 0.9 | -41 |
| 2-EphA1 | -39/3.6 | 1.3 | 3.6/6.2/4.8 | 0.6 | 6.2 | -52 |
| 3-EphA2 | 16/2.1 | 0.6 | 5.5/5.1/5.3 | 0.5 | 0.6 | 17 |
| 4-ErbB1 | -34/3.5 | 1.5 | 3.9/3.6/3.9 | 0.4 | 5.0 | -42 |
| 5-ErbB1/2 | -50/3.8 | 2.1 | 3.3/3.3/2.2 | 0.3 | 3.1 | -51 |
| 6-ErbB2 | -8/1.8 | 4.3 | 4.6/4.1/5.9 | 0.3 | 2.0 | -43 |
| 7-ErbB3 | -26/2.0 | 3.6 | 5.2/4.0/3.3 | 0.4 | 3.0 | 30 |
| 8-ErbB4 | -35/3.8 | 0.5 | 4.1/3.5/5.2 | 0.6 | 1.5 | -41 |
| 9-FGFR3 | 37/3.1 | 0.4 | 6.0/5.1/6.2 | 0.5 | 1.2 | 35 |
| 10-GpA | -43/4.0 | 0.4 | 2.8/4.3/3.8 | 0.6 | 1.0 | -39 |
| 11-PDGFRb | 25/2.1 | 0.6 | 5.8/7.6/5.3 | 0.6 | 0.9 | 23 |

**Table S5:** Comparison of the contact regions of the TM dimers obtained from CG simulation with the NMR structures. Five representative structures from each population cluster are considered for analysis. Contact residues are shown in red. Only backbone contacts are shown here, as detected by PREDDIMER.

| 1st population | 2nd population | 3rd population | NMR |
|---|---|---|---|
| **1-Bnip3** | | | |
| VFLPSLLLSHLLAIGLGIYIG<br>VFLPSLLLSHLLAIGLGIYIG | VFLPSLLLSHLLAIGLGIYIG<br>VFLPSLLLSHLLAIGLGIYIG | VFLPSLLLSHLLAIGLGIYIG<br>VFLPSLLLSHLLAIGLGIYIG | VFLPSLLLSHLLAIGLGIYIG<br>VFLPSLLLSHLLAIGLGIYIG |
| VFLPSLLLSHLLAIGLGIYIG<br>VFLPSLLLSHLLAIGLGIYIG | VFLPSLLLSHLLAIGLGIYIG<br>VFLPSLLLSHLLAIGLGIYIG | VFLPSLLLSHLLAIGLGIYIG<br>VFLPSLLLSHLLAIGLGIYIG | |
| VFLPSLLLSHLLAIGLGIYIG<br>VFLPSLLLSHLLAIGLGIYIG | VFLPSLLLSHLLAIGLGIYIG<br>VFLPSLLLSHLLAIGLGIYIG | VFLPSLLLSHLLAIGLGIYIG<br>VFLPSLLLSHLLAIGLGIYIG | |
| VFLPSLLLSHLLAIGLGIYIG<br>VFLPSLLLSHLLAIGLGIYIG | VFLPSLLLSHLLAIGLGIYIG<br>VFLPSLLLSHLLAIGLGIYIG | VFLPSLLLSHLLAIGLGIYIG<br>VFLPSLLLSHLLAIGLGIYIG | |
| VFLPSLLLSHLLAIGLGIYIG<br>VFLPSLLLSHLLAIGLGIYIG | VFLPSLLLSHLLAIGLGIYIG<br>VFLPSLLLSHLLAIGLGIYIG | VFLPSLLLSHLLAIGLGIYIG<br>VFLPSLLLSHLLAIGLGIYIG | |
| **2-EphA1** | | | |
| IVAVIFGLLLGAALLLGILVF<br>IVAVIFGLLLGAALLLGILVF | IVAVIFGLLLGAALLLGILVF<br>IVAVIFGLLLGAALLLGILVF | IVAVIFGLLLGAALLLGILVF<br>IVAVIFGLLLGAALLLGILVF | IVAVIFGLLLGAALLLGILVF<br>IVAVIFGLLLGAALLLGILVF |
| IVAVIFGLLLGAALLLGILVF<br>IVAVIFGLLLGAALLLGILVF | IVAVIFGLLLGAALLLGILVF<br>IVAVIFGLLLGAALLLGILVF | IVAVIFGLLLGAALLLGILVF<br>IVAVIFGLLLGAALLLGILVF | |
| IVAVIFGLLLGAALLLGILVF<br>IVAVIFGLLLGAALLLGILVF | IVAVIFGLLLGAALLLGILVF<br>IVAVIFGLLLGAALLLGILVF | IVAVIFGLLLGAALLLGILVF<br>IVAVIFGLLLGAALLLGILVF | |
| IVAVIFGLLLGAALLLGILVF<br>IVAVIFGLLLGAALLLGILVF | IVAVIFGLLLGAALLLGILVF<br>IVAVIFGLLLGAALLLGILVF | IVAVIFGLLLGAALLLGILVF<br>IVAVIFGLLLGAALLLGILVF | |
| IVAVIFGLLLGAALLLGILVF<br>IVAVIFGLLLGAALLLGILVF | IVAVIFGLLLGAALLLGILVF<br>IVAVIFGLLLGAALLLGILVF | IVAVIFGLLLGAALLLGILVF<br>IVAVIFGLLLGAALLLGILVF | |
| **3-EphA2** | | | |
| IGGVAVGVVLLLVLAGVGFFI<br>IGGVAVGVVLLLVLAGVGFFI | IGGVAVGVVLLLVLAGVGFFI<br>IGGVAVGVVLLLVLAGVGFFI | IGGVAVGVVLLLVLAGVGFFI<br>IGGVAVGVVLLLVLAGVGFFI | IGGVAVGVVLLLVLAGVGFFI<br>IGGVAVGVVLLLVLAGVGFFI |
| IGGVAVGVVLLLVLAGVGFFI<br>IGGVAVGVVLLLVLAGVGFFI | IGGVAVGVVLLLVLAGVGFFI<br>IGGVAVGVVLLLVLAGVGFFI | IGGVAVGVVLLLVLAGVGFFI<br>IGGVAVGVVLLLVLAGVGFFI | |
| IGGVAVGVVLLLVLAGVGFFI<br>IGGVAVGVVLLLVLAGVGFFI | IGGVAVGVVLLLVLAGVGFFI<br>IGGVAVGVVLLLVLAGVGFFI | IGGVAVGVVLLLVLAGVGFFI<br>IGGVAVGVVLLLVLAGVGFFI | |
| IGGVAVGVVLLLVLAGVGFFI<br>IGGVAVGVVLLLVLAGVGFFI | IGGVAVGVVLLLVLAGVGFFI<br>IGGVAVGVVLLLVLAGVGFFI | IGGVAVGVVLLLVLAGVGFFI<br>IGGVAVGVVLLLVLAGVGFFI | |
| IGGVAVGVVLLLVLAGVGFFI<br>IGGVAVGVVLLLVLAGVGFFI | IGGVAVGVVLLLVLAGVGFFI<br>IGGVAVGVVLLLVLAGVGFFI | IGGVAVGVVLLLVLAGVGFFI<br>IGGVAVGVVLLLVLAGVGFFI | |
| **4-ErbB1** | | | |
| IATGMVGALLLLLVVALGIGLFM<br>IATGMVGALLLLLVVALGIGLFM | IATGMVGALLLLLVVALGIGLFM<br>IATGMVGALLLLLVVALGIGLFM | IATGMVGALLLLLVVALGIGLFM<br>IATGMVGALLLLLVVALGIGLFM | IATGMVGALLLLLVVALGIGLFM<br>IATGMVGALLLLLVVALGIGLFM |

| | | | |
|---|---|---|---|
| IATGMVGALLLLLVVALGIGLFM<br>IATGMVGALLLLLVVALGIGLFM | IATGMVGALLLLLVVALGIGLFM<br>IATGMVGALLLLLVVALGIGLFM | IATGMVGALLLLLVVALGIGLFM<br>IATGMVGALLLLLVVALGIGLFM | |
| IATGMVGALLLLLVVALGIGLFM<br>IATGMVGALLLLLVVALGIGLFM | IATGMVGALLLLLVVALGIGLFM<br>IATGMVGALLLLLVVALGIGLFM | IATGMVGALLLLLVVALGIGLFM<br>IATGMVGALLLLLVVALGIGLFM | |
| IATGMVGALLLLLVVALGIGLFM<br>IATGMVGALLLLLVVALGIGLFM | IATGMVGALLLLLVVALGIGLFM<br>IATGMVGALLLLLVVALGIGLFM | IATGMVGALLLLLVVALGIGLFM<br>IATGMVGALLLLLVVALGIGLFM | |
| IATGMVGALLLLLVVALGIGLFM<br>IATGMVGALLLLLVVALGIGLFM | IATGMVGALLLLLVVALGIGLFM<br>IATGMVGALLLLLVVALGIGLFM | IATGMVGALLLLLVVALGIGLFM<br>IATGMVGALLLLLVVALGIGLFM | |
| **5-ErbB1-B2** | | | |
| IATGMVGALLLLLVVALGIGLFM<br>SIISAVVGILLVVVLGVVFGILI | IATGMVGALLLLLVVALGIGLFM<br>SIISAVVGILLVVVLGVVFGILI | IATGMVGALLLLLVVALGIGLFM<br>SIISAVVGILLVVVLGVVFGILI | IATGMVGALLLLLVVALGIGLFM<br>SIISAVVGILLVVVLGVVFGILI |
| IATGMVGALLLLLVVALGIGLFM<br>SIISAVVGILLVVVLGVVFGILI | IATGMVGALLLLLVVALGIGLFM<br>SIISAVVGILLVVVLGVVFGILI | IATGMVGALLLLLVVALGIGLFM<br>SIISAVVGILLVVVLGVVFGILI | |
| IATGMVGALLLLLVVALGIGLFM<br>SIISAVVGILLVVVLGVVFGILI | IATGMVGALLLLLVVALGIGLFM<br>SIISAVVGILLVVVLGVVFGILI | IATGMVGALLLLLVVALGIGLFM<br>SIISAVVGILLVVVLGVVFGILI | |
| IATGMVGALLLLLVVALGIGLFM<br>SIISAVVGILLVVVLGVVFGILI | IATGMVGALLLLLVVALGIGLFM<br>SIISAVVGILLVVVLGVVFGILI | IATGMVGALLLLLVVALGIGLFM<br>SIISAVVGILLVVVLGVVFGILI | |
| IATGMVGALLLLLVVALGIGLFM<br>SIISAVVGILLVVVLGVVFGILI | IATGMVGALLLLLVVALGIGLFM<br>SIISAVVGILLVVVLGVVFGILI | IATGMVGALLLLLVVALGIGLFM<br>SIISAVVGILLVVVLGVVFGILI | |
| **6-ErbB2** | | | |
| SIISAVVGILLVVVLGVVFGILI<br>SIISAVVGILLVVVLGVVFGILI | SIISAVVGILLVVVLGVVFGILI<br>SIISAVVGILLVVVLGVVFGILI | SIISAVVGILLVVVLGVVFGILI<br>SIISAVVGILLVVVLGVVFGILI | SIISAVVGILLVVVLGVVFGILI<br>SIISAVVGILLVVVLGVVFGILI |
| SIISAVVGILLVVVLGVVFGILI<br>SIISAVVGILLVVVLGVVFGILI | SIISAVVGILLVVVLGVVFGILI<br>SIISAVVGILLVVVLGVVFGILI | SIISAVVGILLVVVLGVVFGILI<br>SIISAVVGILLVVVLGVVFGILI | |
| SIISAVVGILLVVVLGVVFGILI<br>SIISAVVGILLVVVLGVVFGILI | SIISAVVGILLVVVLGVVFGILI<br>SIISAVVGILLVVVLGVVFGILI | SIISAVVGILLVVVLGVVFGILI<br>SIISAVVGILLVVVLGVVFGILI | |
| SIISAVVGILLVVVLGVVFGILI<br>SIISAVVGILLVVVLGVVFGILI | SIISAVVGILLVVVLGVVFGILI<br>SIISAVVGILLVVVLGVVFGILI | SIISAVVGILLVVVLGVVFGILI<br>SIISAVVGILLVVVLGVVFGILI | |
| SIISAVVGILLVVVLGVVFGILI<br>SIISAVVGILLVVVLGVVFGILI | SIISAVVGILLVVVLGVVFGILI<br>SIISAVVGILLVVVLGVVFGILI | SIISAVVGILLVVVLGVVFGILI<br>SIISAVVGILLVVVLGVVFGILI | |
| **7-ErbB3** | | | |
| MALTVIAGLVVIFMMLGGTFL<br>MALTVIAGLVVIFMMLGGTFL | MALTVIAGLVVIFMMLGGTFL<br>MALTVIAGLVVIFMMLGGTFL | MALTVIAGLVVIFMMLGGTFL<br>MALTVIAGLVVIFMMLGGTFL | MALTVIAGLVVIFMMLGGTFL<br>MALTVIAGLVVIFMMLGGTFL |
| MALTVIAGLVVIFMMLGGTFL<br>MALTVIAGLVVIFMMLGGTFL | MALTVIAGLVVIFMMLGGTFL<br>MALTVIAGLVVIFMMLGGTFL | MALTVIAGLVVIFMMLGGTFL<br>MALTVIAGLVVIFMMLGGTFL | |
| MALTVIAGLVVIFMMLGGTFL<br>MALTVIAGLVVIFMMLGGTFL | MALTVIAGLVVIFMMLGGTFL<br>MALTVIAGLVVIFMMLGGTFL | MALTVIAGLVVIFMMLGGTFL<br>MALTVIAGLVVIFMMLGGTFL | |
| MALTVIAGLVVIFMMLGGTFL<br>MALTVIAGLVVIFMMLGGTFL | MALTVIAGLVVIFMMLGGTFL<br>MALTVIAGLVVIFMMLGGTFL | MALTVIAGLVVIFMMLGGTFL<br>MALTVIAGLVVIFMMLGGTFL | |
| MALTVIAGLVVIFMMLGGTFL<br>MALTVIAGLVVIFMMLGGTFL | MALTVIAGLVVIFMMLGGTFL<br>MALTVIAGLVVIFMMLGGTFL | MALTVIAGLVVIFMMLGGTFL<br>MALTVIAGLVVIFMMLGGTFL | |
| **8-ErbB4** | | | |
| LIAAGVIGGLFILVIVGLTFAVYV<br>LIAAGVIGGLFILVIVGLTFAVYV | LIAAGVIGGLFILVIVGLTFAVYV<br>LIAAGVIGGLFILVIVGLTFAVYV | LIAAGVIGGLFILVIVGLTFAVYV<br>LIAAGVIGGLFILVIVGLTFAVYV | LIAAGVIGGLFILVIVGLTFAVYV<br>LIAAGVIGGLFILVIVGLTFAVYV |

| | | | |
|---|---|---|---|
| LIAAGVIGGLFILVIVGLTFAVYV<br>LIAAGVIGGLFILVIVGLTFAVYV | LIAAGVIGGLFILVIVGLTFAVYV<br>LIAAGVIGGLFILVIVGLTFAVYV | LIAAGVIGGLFILVIVGLTFAVYV<br>LIAAGVIGGLFILVIVGLTFAVYV | |
| LIAAGVIGGLFILVIVGLTFAVYV<br>LIAAGVIGGLFILVIVGLTFAVYV | LIAAGVIGGLFILVIVGLTFAVYV<br>LIAAGVIGGLFILVIVGLTFAVYV | LIAAGVIGGLFILVIVGLTFAVYV<br>LIAAGVIGGLFILVIVGLTFAVYV | |
| LIAAGVIGGLFILVIVGLTFAVYV<br>LIAAGVIGGLFILVIVGLTFAVYV | LIAAGVIGGLFILVIVGLTFAVYV<br>LIAAGVIGGLFILVIVGLTFAVYV | LIAAGVIGGLFILVIVGLTFAVYV<br>LIAAGVIGGLFILVIVGLTFAVYV | |
| LIAAGVIGGLFILVIVGLTFAVYV<br>LIAAGVIGGLFILVIVGLTFAVYV | LIAAGVIGGLFILVIVGLTFAVYV<br>LIAAGVIGGLFILVIVGLTFAVYV | LIAAGVIGGLFILVIVGLTFAVYV<br>LIAAGVIGGLFILVIVGLTFAVYV | |
| **9-FGFR3** | | | |
| ILSYGVGFFLFILVVAAVTLC<br>ILSYGVGFFLFILVVAAVTLC | ILSYGVGFFLFILVVAAVTLC<br>ILSYGVGFFLFILVVAAVTLC | ILSYGVGFFLFILVVAAVTLC<br>ILSYGVGFFLFILVVAAVTLC | ILSYGVGFFLFILVVAAVTLC<br>ILSYGVGFFLFILVVAAVTLC |
| ILSYGVGFFLFILVVAAVTLC<br>ILSYGVGFFLFILVVAAVTLC | ILSYGVGFFLFILVVAAVTLC<br>ILSYGVGFFLFILVVAAVTLC | ILSYGVGFFLFILVVAAVTLC<br>ILSYGVGFFLFILVVAAVTLC | |
| ILSYGVGFFLFILVVAAVTLC<br>ILSYGVGFFLFILVVAAVTLC | ILSYGVGFFLFILVVAAVTLC<br>ILSYGVGFFLFILVVAAVTLC | ILSYGVGFFLFILVVAAVTLC<br>ILSYGVGFFLFILVVAAVTLC | |
| ILSYGVGFFLFILVVAAVTLC<br>ILSYGVGFFLFILVVAAVTLC | ILSYGVGFFLFILVVAAVTLC<br>ILSYGVGFFLFILVVAAVTLC | ILSYGVGFFLFILVVAAVTLC<br>ILSYGVGFFLFILVVAAVTLC | |
| ILSYGVGFFLFILVVAAVTLC<br>ILSYGVGFFLFILVVAAVTLC | ILSYGVGFFLFILVVAAVTLC<br>ILSYGVGFFLFILVVAAVTLC | ILSYGVGFFLFILVVAAVTLC<br>ILSYGVGFFLFILVVAAVTLC | |
| **10-GpA** | | | |
| ITLIIFGVMAGVIGTILLISYGI<br>ITLIIFGVMAGVIGTILLISYGI | ITLIIFGVMAGVIGTILLISYGI<br>ITLIIFGVMAGVIGTILLISYGI | ITLIIFGVMAGVIGTILLISYGI<br>ITLIIFGVMAGVIGTILLISYGI | ITLIIFGVMAGVIGTILLISYGI<br>ITLIIFGVMAGVIGTILLISYGI |
| ITLIIFGVMAGVIGTILLISYGI<br>ITLIIFGVMAGVIGTILLISYGI | ITLIIFGVMAGVIGTILLISYGI<br>ITLIIFGVMAGVIGTILLISYGI | ITLIIFGVMAGVIGTILLISYGI<br>ITLIIFGVMAGVIGTILLISYGI | |
| ITLIIFGVMAGVIGTILLISYGI<br>ITLIIFGVMAGVIGTILLISYGI | ITLIIFGVMAGVIGTILLISYGI<br>ITLIIFGVMAGVIGTILLISYGI | ITLIIFGVMAGVIGTILLISYGI<br>ITLIIFGVMAGVIGTILLISYGI | |
| ITLIIFGVMAGVIGTILLISYGI<br>ITLIIFGVMAGVIGTILLISYGI | ITLIIFGVMAGVIGTILLISYGI<br>ITLIIFGVMAGVIGTILLISYGI | ITLIIFGVMAGVIGTILLISYGI<br>ITLIIFGVMAGVIGTILLISYGI | |
| ITLIIFGVMAGVIGTILLISYGI<br>ITLIIFGVMAGVIGTILLISYGI | ITLIIFGVMAGVIGTILLISYGI<br>ITLIIFGVMAGVIGTILLISYGI | ITLIIFGVMAGVIGTILLISYGI<br>ITLIIFGVMAGVIGTILLISYGI | |
| **11-PDGFRb** | | | |
| VVISAILALVVLTIISLIILI<br>VVISAILALVVLTIISLIILI | VVISAILALVVLTIISLIILI<br>VVISAILALVVLTIISLIILI | VVISAILALVVLTIISLIILI<br>VVISAILALVVLTIISLIILI | VVISAILALVVLTIISLIILI<br>VVISAILALVVLTIISLIILI |
| VVISAILALVVLTIISLIILI<br>VVISAILALVVLTIISLIILI | VVISAILALVVLTIISLIILI<br>VVISAILALVVLTIISLIILI | VVISAILALVVLTIISLIILI<br>VVISAILALVVLTIISLIILI | |
| VVISAILALVVLTIISLIILI<br>VVISAILALVVLTIISLIILI | VVISAILALVVLTIISLIILI<br>VVISAILALVVLTIISLIILI | VVISAILALVVLTIISLIILI<br>VVISAILALVVLTIISLIILI | |
| VVISAILALVVLTIISLIILI<br>VVISAILALVVLTIISLIILI | VVISAILALVVLTIISLIILI<br>VVISAILALVVLTIISLIILI | VVISAILALVVLTIISLIILI<br>VVISAILALVVLTIISLIILI | |
| VVISAILALVVLTIISLIILI<br>VVISAILALVVLTIISLIILI | VVISAILALVVLTIISLIILI<br>VVISAILALVVLTIISLIILI | VVISAILALVVLTIISLIILI<br>VVISAILALVVLTIISLIILI | |

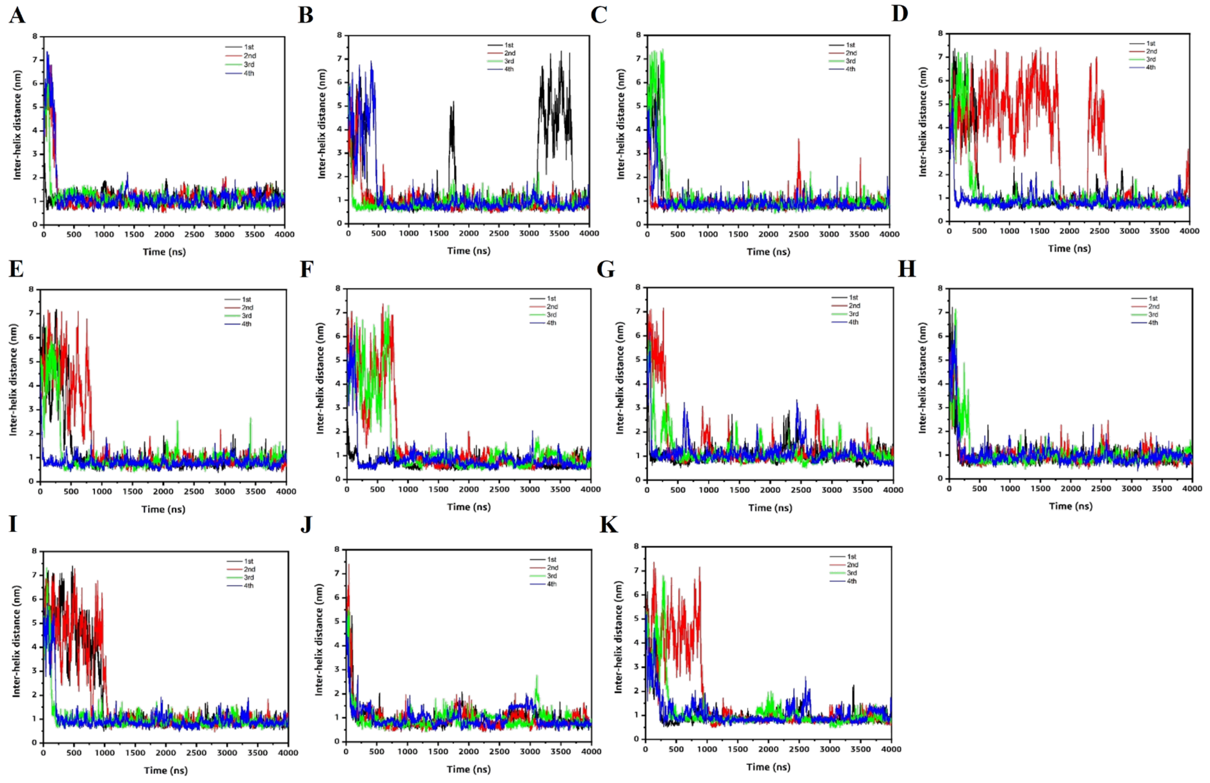


**Fig. S1** Comparison of inter-helix distance between the TM monomers as a function of simulation time. (A) Bnip3 (B) EphA1 (C) EphA2 (D) ErbB1 (E) ErbB1/2 (F) ErbB2 (G) ErbB3 (H) ErbB4 (I) FGFR3 (J) GpA and (K) PDGFRb. 1st, 2nd, 3rd and 4th simulation results are shown black, red, green and blue lines.

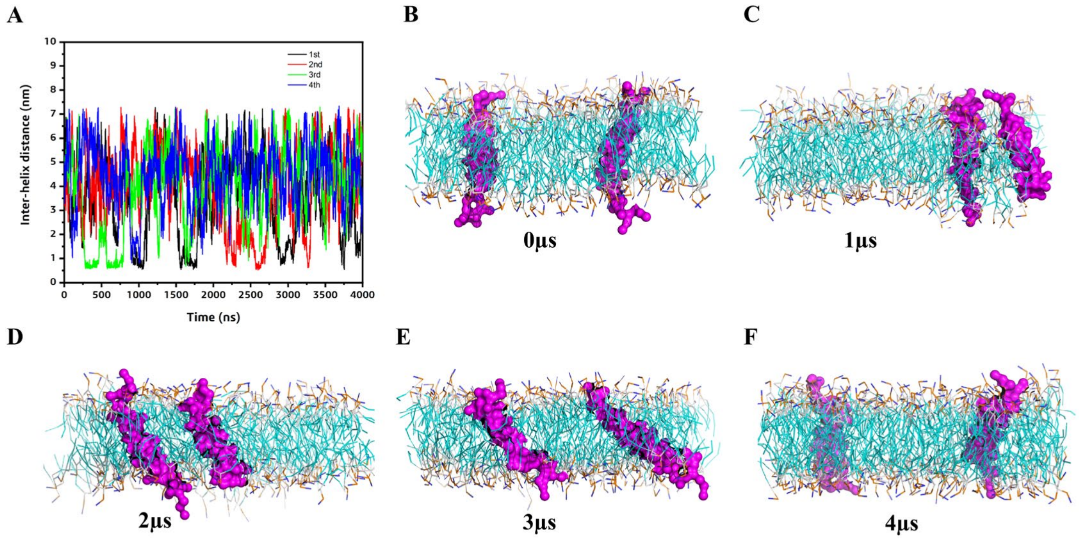


**Fig. S2 Control simulation of polyLeu ($K_2$-$L_{26}$-$K_2$) peptides in DMPC bilayer.** (A) Comparison of inter-helix distance between the TM monomers as a function of simulation time. 1st, 2nd, 3rd and 4th simulation results are shown black, red, green and blue lines. (B-F) Snapshots of the peptides (shown in magenta) at different time frames suggesting no interaction between them over 4 μs.

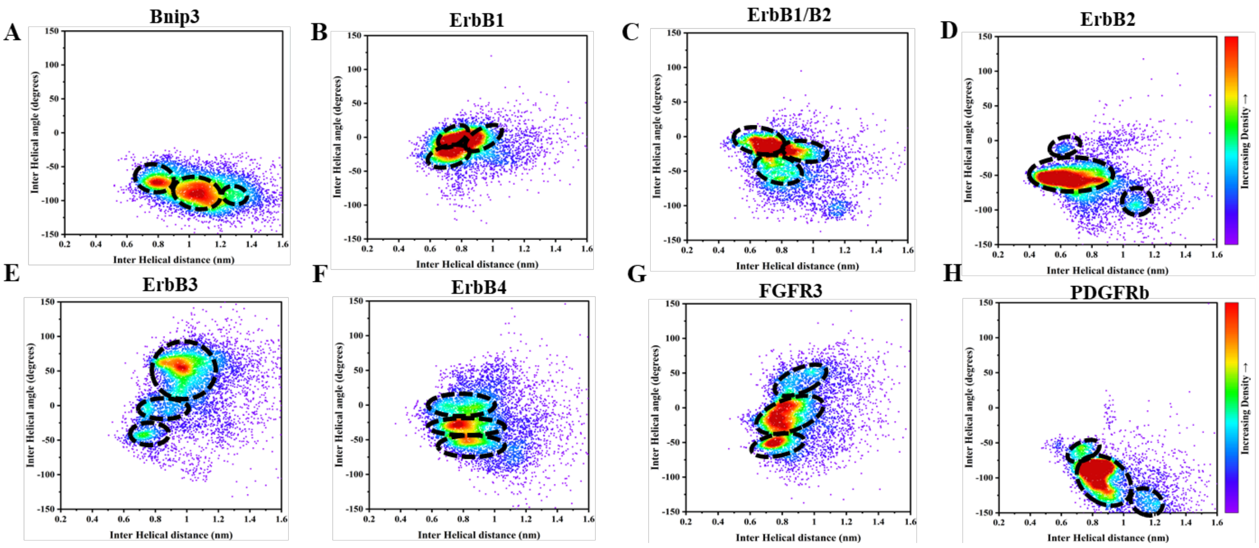


**Fig. S3** 2D distribution plot (interhelix angle vs. distance) for (A) Bnip3 (B) ErbB1 (C) ErbB1/2 (D) ErbB2 (E) ErbB3 (F) ErbB4 (G) FGFR3 and (H) PDGFRb. Each plot is divided into 3 distinct population clusters based on the inter-helical angle and the inter-helical distance. The population of each cluster are shown in Table1. Several structures from each population cluster was extracted and then compared with the NMR structure (shown in Table1). Data from the last 2.5μs simulations are considered. Data points at an interval of 500ps are skipped for clarity.

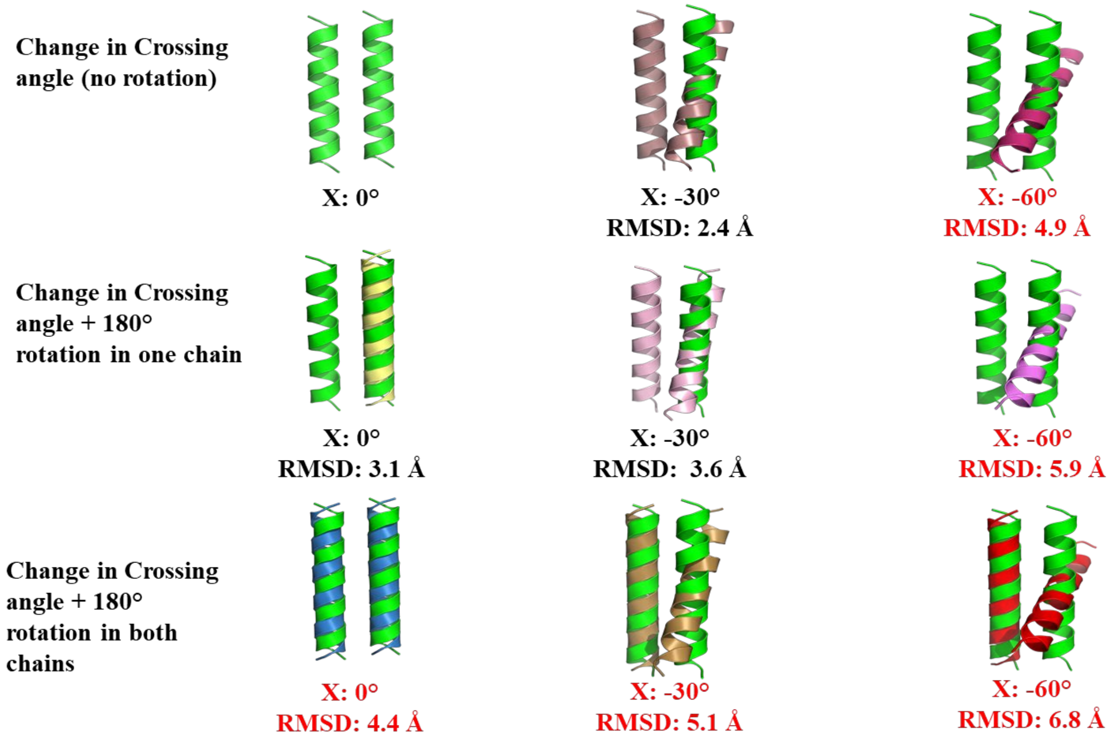


**Fig. S4** Example of RMSD reference for changes for an ideal parallel TM dimer. The modeled dimer of EphA1 shows variation in backbone RMSD compared to the starting parallel conformation (green), when there is change in crossing angle (X) and either rotation (180° shown here) in either one or both the chains. RMSD range > 4.5 Å are marked as red.

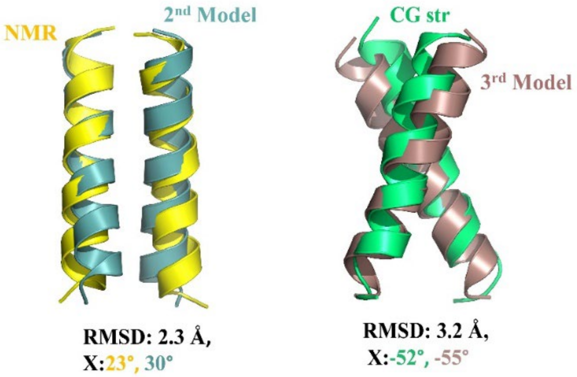


**Fig. S5** Superimposition of the best fit PREDDIMER models onto the best fit CG simulated structure (shown in green) and onto the NMR structure (shown in yellow) for PDGFRb.

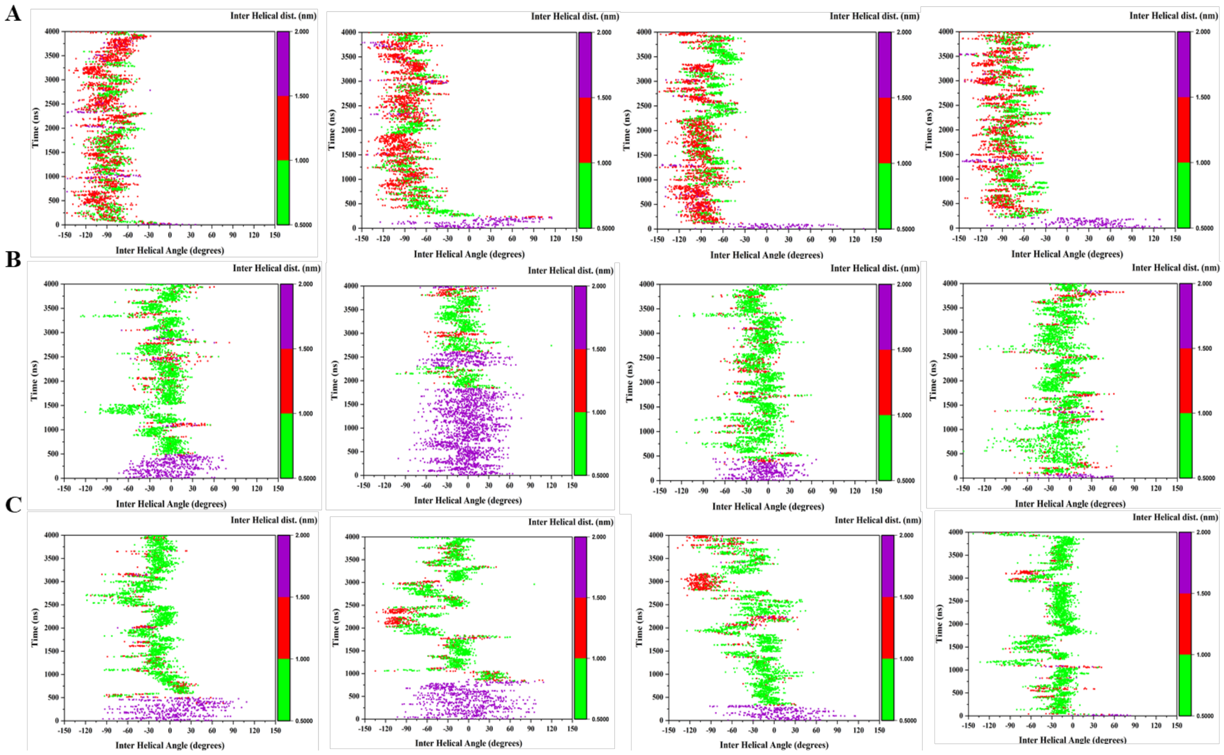


**Fig. S6 2D Plots showing the conformational transition of TM dimers over the simulation time considering the inter-helical angle vs inter-helical distance for (A) Bnip3, (B) ErbB1 and (C) ErbB1/ B2.** Results from 4 trajectories are shown here. The plots are colored based on the inter-helical distance from 0.5- 1.0 nm (green), 1- 1.5 nm (red) and 1.5- 2.0 nm (purple).

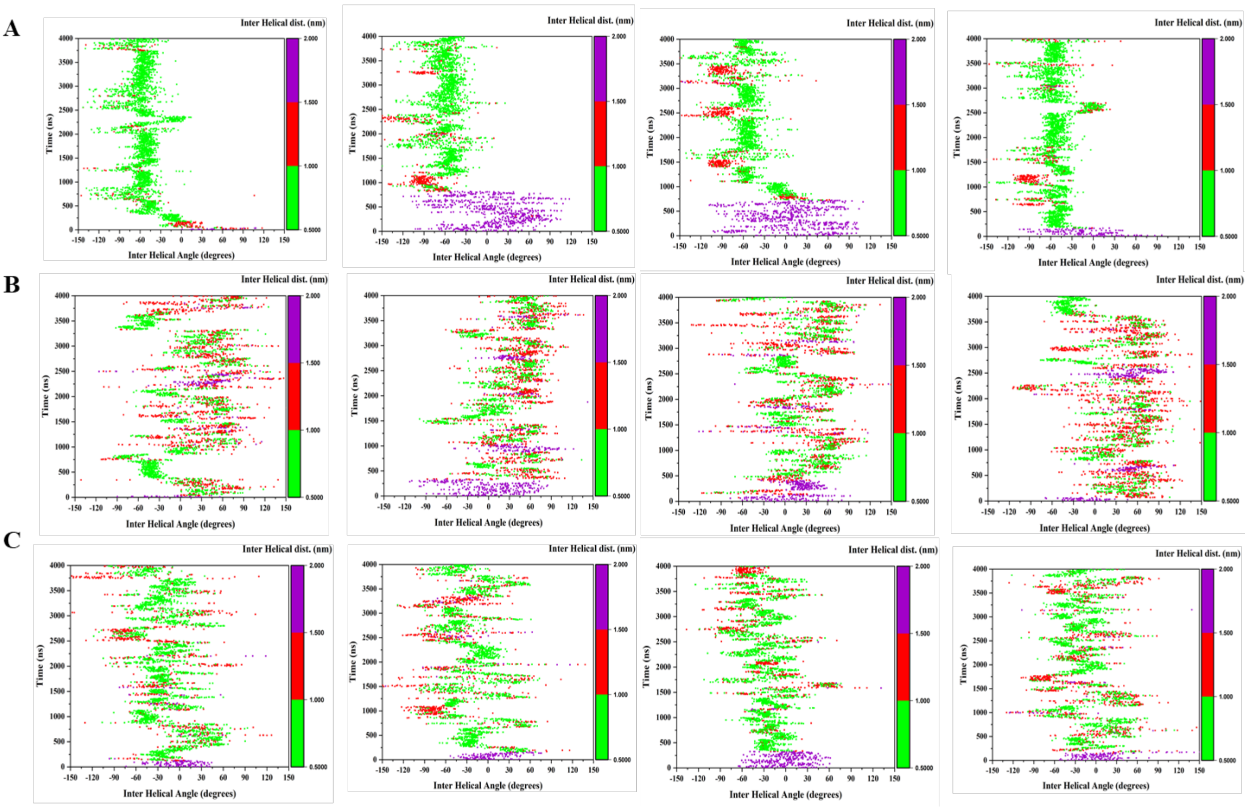


**Fig. S7 2D Plots showing the conformational transition of TM dimers over the simulation time considering the inter-helical angle vs inter-helical distance for (A) ErbB2, (B) ErbB3 and (C) ErbB4.** Results from 4 trajectories are shown here. The plots are colored based on the inter-helical distance from 0.5- 1.0 nm (green), 1- 1.5 nm (red) and 1.5- 2.0 nm (purple).

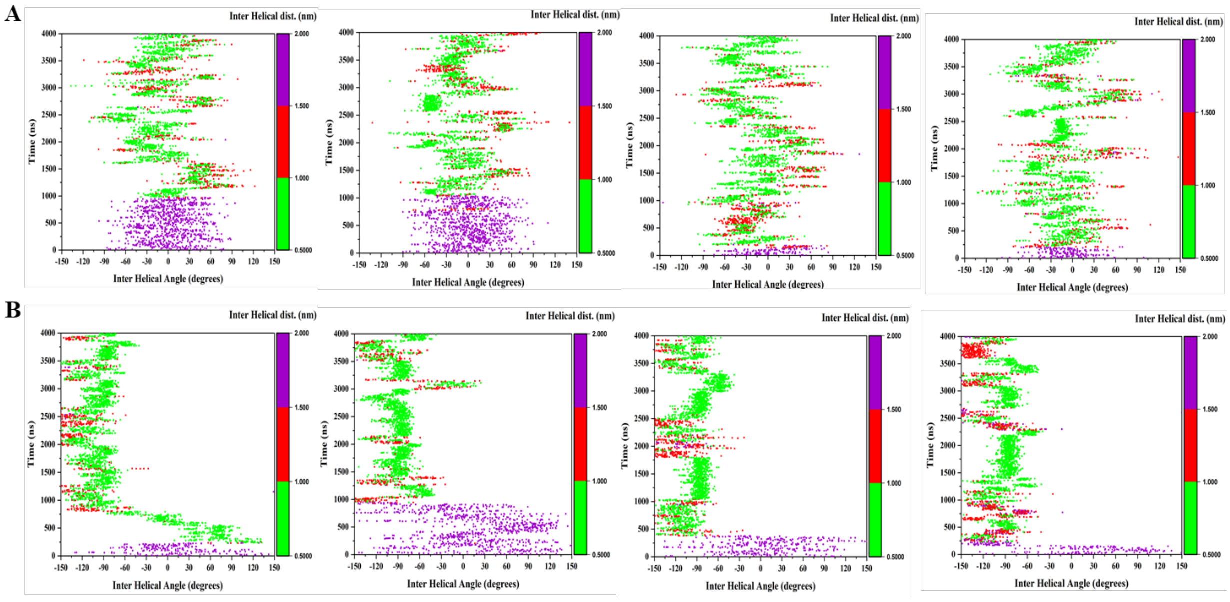


**Fig. S8 2D Plots showing the conformational transition of TM dimers over the simulation time considering the inter-helical angle vs inter-helical distance for (A) FGFR3 and (B)**

**PDGFRb.** Results from 4 trajectories are shown here. The plots are colored based on the inter-helical distance from 0.5- 1.0 nm (green), 1- 1.5 nm (red) and 1.5- 2.0 nm (purple).

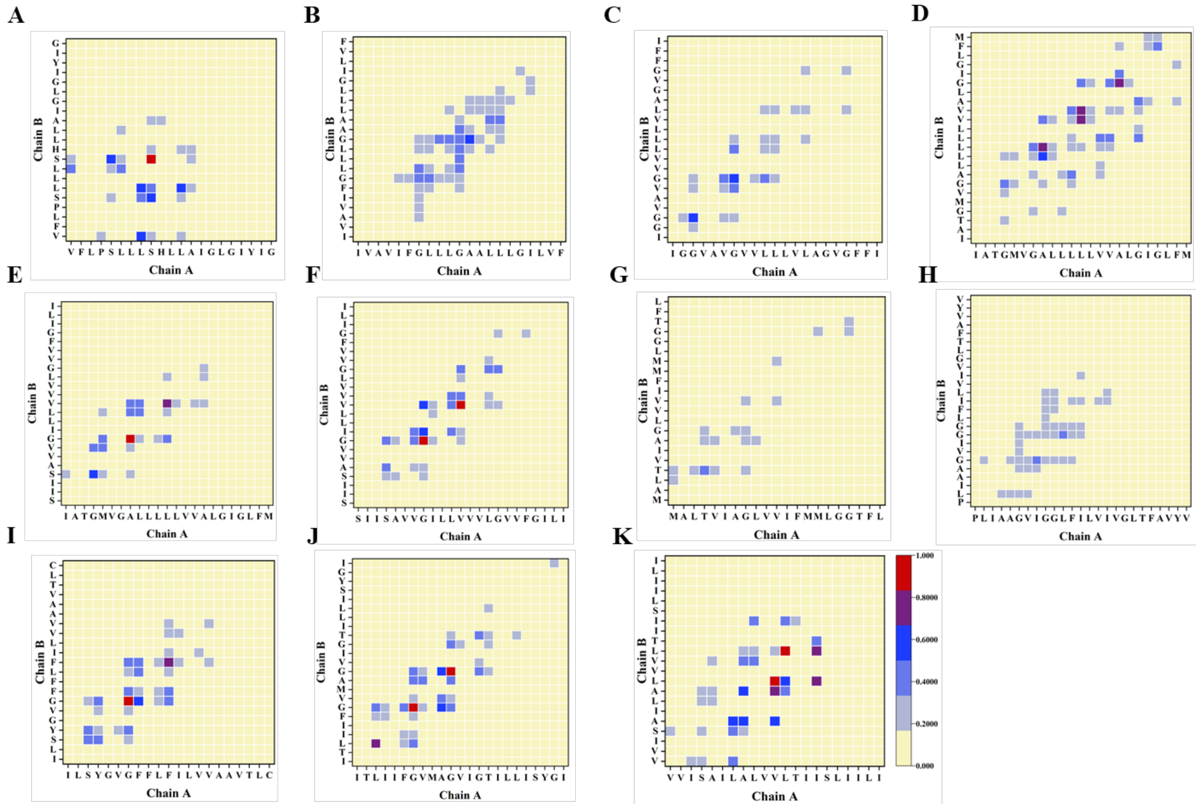


**Fig. S9 Comparison of contact map interface between the mainchain of helices in the dimers of all 11 receptor TM regions.** (A) Bnip3 (B) EphA1 (C) EphA2 (D) ErbB1 (E) ErbB1/2 (F) ErbB2 (G) ErbB3 (H) ErbB4 (I) FGFR3 (J) GpA and (K) PDGFRb.

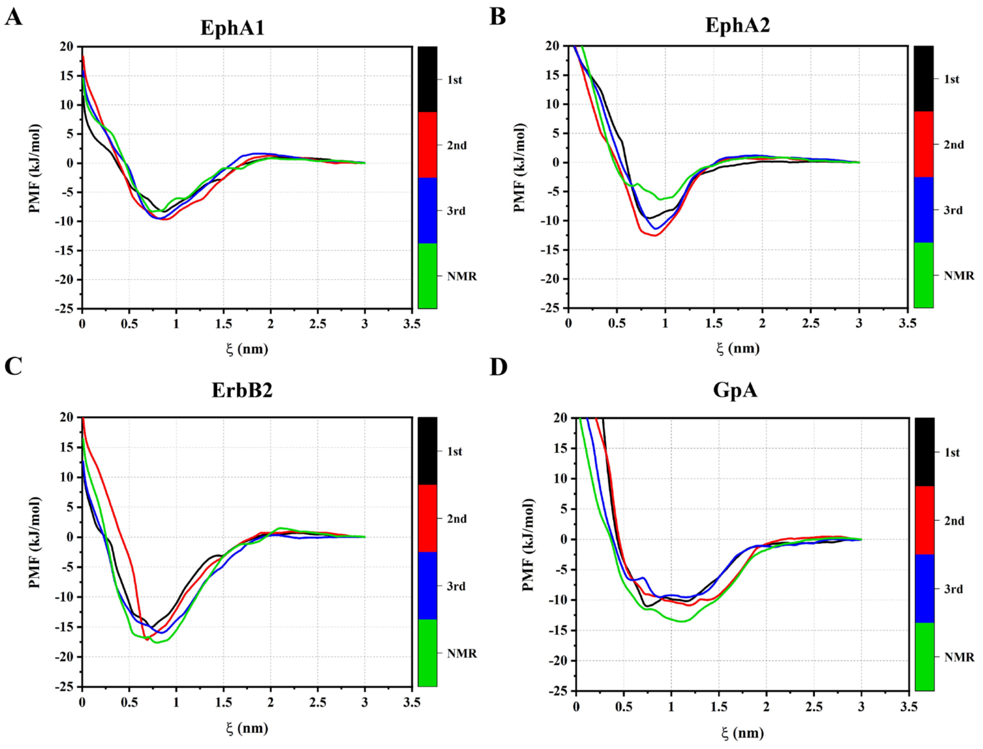


**Fig. S10 PMF for dimer association of EphA1, EphA2, ErbB2 and GpA** as a function of distance between the center of mass of TM peptides for the best three structures (1$^{st}$, 2$^{nd}$ and 3$^{rd}$) among the three population clusters. Additional PMFs for the NMR conformations are represented as green lines. PMF values for the 1$^{st}$, 2$^{nd}$ and 3$^{rd}$ configurations are shown as black, red and blue lines.

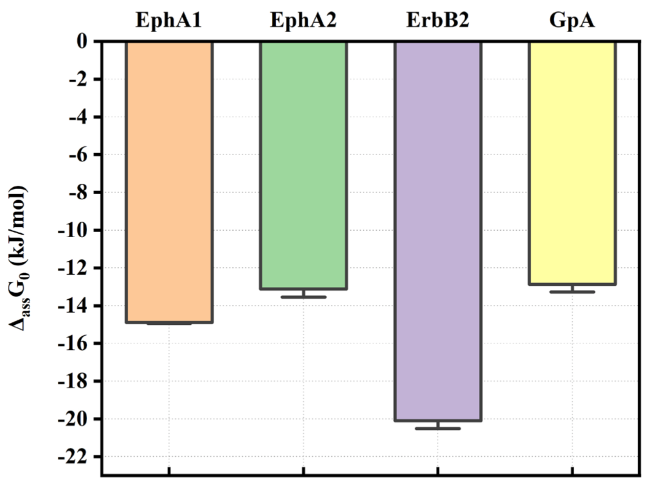


**Fig. S11 Comparison of average free energy ($\Delta_{ass}G_0$) of TM dimerization for EphA1, EphA2, ErbB2 and GpA.**

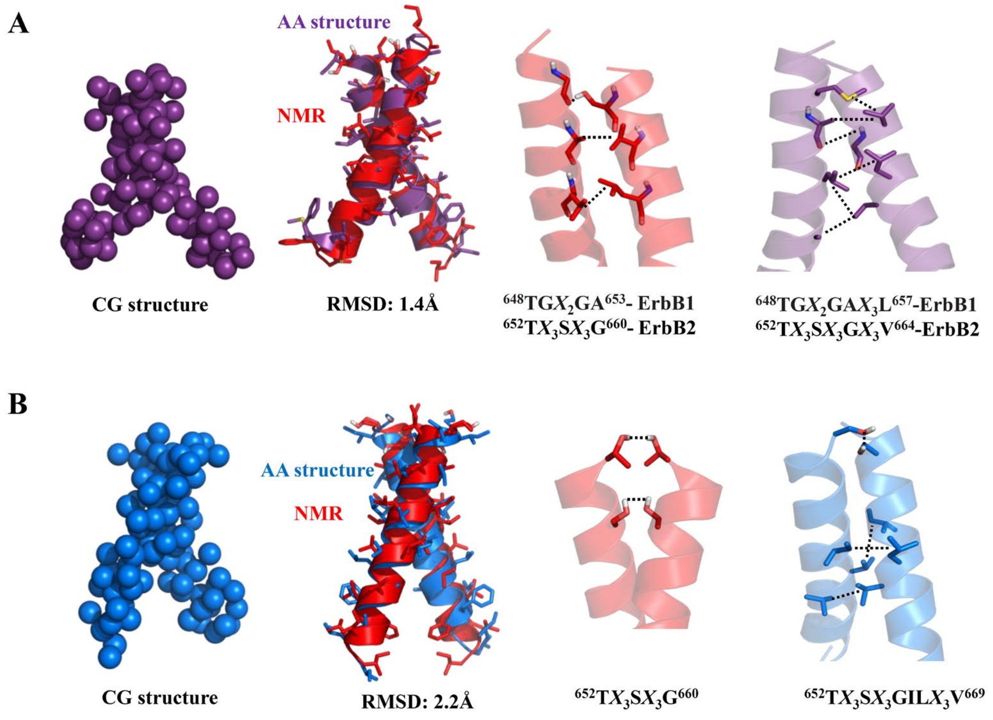


**Fig. S12** Representative structures of CG and AA configuration and comparison of AA structures with the NMR structures of (A) ErbB1/2 and (B) ErbB2. Interaction motifs of NMR and the simulation structures are compared. All non-bonded interactions are shown as dotted lines.
